# Accumulation of LRRK2-associated phospho-Rab12 degenerative lysosomes in tauopathies

**DOI:** 10.1101/2025.06.06.658328

**Authors:** Silas A. Buck, Tuyana Malankhanova, Eileen B. Ma, Sarah Yim, Harrison W. Pratt, John Ervin, Shih-Hsiu J. Wang, Todd J. Cohen, Andrew B. West, Laurie H. Sanders

## Abstract

Parkinson’s disease (PD) pathogenic mutations in *leucine-rich repeat kinase 2* (*LRRK2*) are associated with endolysosomal dysfunction across cell types, and carriers of *LRRK2* mutations variably present with phosphorylated tau and α-synuclein deposits in post-mortem analysis. *LRRK2* mutations increase the phosphorylation of Rab substrates including Rab12. Rab12 is expressed in neuronal and non-neuronal cells with localization to membranes in the endolysosomal compartment. Under lysosomal stress, LRRK2 interaction with Rab12 upregulates LRRK2 kinase activity. In this study, using a recently developed monoclonal antibody directed to the LRRK2-mediated phosphorylation site on Rab12 at amino acid Ser106 (pS106-Rab12), we test whether aberrant LRRK2 phosphorylation is associated with tau and/or α-synuclein pathology across clinically distinct neurodegenerative diseases. Analysis of brain tissue lysates and immunohistochemistry of pathology-susceptible brain regions demonstrate that pS106-Rab12 levels are increased in Dementia with Lewy bodies (DLB), Alzheimer’s disease (AD), and PD, and in *LRRK2* mutation carriers. In early pathological stages, phosphorylated Rab12 localizes to granulovacuolar degeneration bodies (GVBs), which are thought to be active lysosomal-like structures, in neurons. pS106-Rab12-positive GVBs accumulate with pathological tau across brain tissues in DLB, AD, and PD, and in *LRRK2* mutation carriers. In a mouse model of tauopathy, pS106-Rab12 localizes to GVBs during early tau deposition in an age-dependent manner. While GVBs are largely absent in neurons with mature protein pathology, subsets of both tau and α-synuclein inclusions appear to incorporate pS106-Rab12 at later pathological stages. Finally, pS106-Rab12 labels GVBs in neurons and shows widespread co-pathology with tau inclusions in primary tauopathies including Pick’s disease, progressive supranuclear palsy and corticobasal degeneration. These results implicate LRRK2 kinase activity and Rab phosphorylation in endolysosomal dysfunction in both tau and α-synuclein-associated neurodegenerative diseases.

## Introduction

Mutations in *leucine-rich repeat kinase 2* (*LRRK2*) increase its kinase activity and cause autosomal dominant Parkinson’s disease (PD) that clinically overlaps with idiopathic PD (iPD), but with pleiotropic pathologies [5, 26, 29, 30, 40, 63, 64, 74, 78, 83, 93, 99]. Dementia associated with tau pathology has been reported in *LRRK2* carriers [6, 32, 38, 43, 70, 86, 99]. About half of LRRK2-associated neurodegenerative disease displays Lewy body pathology and/or Alzheimer’s disease (AD)-associated 3R/4R tau pathology through much of the cortex [6, 32, 43]. LRRK2 is linked with tau pathology by the association of genetic variation at the *LRRK2* locus with survival in the primary tauopathy progressive supranuclear palsy (PSP) [39]. Additionally, some *LRRK2* PD patients display PSP-like symptoms together with hyperphosphorylated tau pathology found on post-mortem immunohistochemistry (IHC) analysis [6, 79, 99]. Consistent with variable α-synuclein and tau co-pathology across LRRK2 PD cases, pathogenic *LRRK2* mutations have been shown to exacerbate both α-synuclein and tau aggregates in some animal models. Rodent models with pathogenic PD-linked *LRRK2* mutations display increased accumulation of insoluble α-synuclein inclusions and greater α-synuclein-induced neurodegeneration, which can be reversed by LRRK2 kinase inhibition [18, 71, 89]. However, other studies show that α-synuclein inclusion formation is independent of pathogenic LRRK2 [23, 24]. A better mechanistic understanding of how LRRK2 activity may mediate initial pathologies, and their later progression, may help resolve discrepancies in the literature. For tau, mice expressing PD-linked pathogenic *LRRK2* mutations display increased accumulation and cell-to-cell transmission of hyperphosphorylated tau [2, 52, 56, 60]. However, other studies suggest no significant effect of *LRRK2* mutations on tau aggregation, depending on the model [24, 57]. These findings highlight discrepancies in our understanding of LRRK2 activity at different stages of disease and how LRRK2 activity might impact α-synuclein and tau pathology.

In cell culture models, LRRK2 is recruited to damaged lysosomes and phosphorylates Rab proteins, including Rab10 and Rab12 [1, 3, 4, 7, 8, 45, 80, 81]. Rab12 interaction with LRRK2 promotes LRRK2-mediated Rab10 phosphorylation [19, 91]. Further, Rab12 regulates lysosomal damage-mediated LRRK2 localization to lysosomes, thereby suggesting Rab12 is both a substrate and regulator of LRRK2 [91]. Considering both pathological α-synuclein and tau accumulation induce endolysosomal stress, and both proteins can be degraded by lysosomes, LRRK2 and Rab12 may play fundamental roles in mediating early stages of α-synuclein and tau pathology in neurodegenerative diseases [12, 34, 41, 50, 54, 66, 68, 69, 77].

In addition to endolysosomal stress, both α-synuclein and tau pathology induce neuropathology associated with the lysosome, named granulovacuolar degeneration bodies (GVBs). GVBs describe the appearance of membrane-delineated structures with a dense granule that accumulate predominantly in neurons of patients with neurodegenerative diseases [94]. GVBs have been described in primary and secondary tauopathies including AD, Pick’s disease, PSP, and corticobasal degeneration (CBD), as well as synucleinopathies like PD and Dementia with Lewy bodies (DLB) [94]. GVBs are proteolytically active lysosomal structures in neuron models (i.e., likely functional lysosomes) that are directly induced by intracellular tau or α-synuclein aggregation, but not amyloid-β deposition [36, 42, 95]. In human tauopathies and mouse tauopathy models, GVBs closely follow the appearance of tau pathology [48, 82, 94]. Intriguingly, the presence of GVBs is associated with early, but not late, stages of pathological tau and α-synuclein inclusion accumulation in neurons [42, 48]. GVBs are mainly restricted to neurons, although some, but not all, investigations report GVBs in glia of Pick’s disease, PSP and CBD cases [61, 72, 87]. Overall, these studies suggest a connection between lysosomal dysfunction, GVBs, and early neuronal tau and α-synuclein aggregation. Indeed, iPD cases display GVBs, and one qualitative study suggests that LRRK2 PD may display enrichment of GVBs compared to iPD cases [35, 55, 82, 94].

In this study, we tested whether aberrant LRRK2 activity is associated with tau and/or α-synuclein pathology in clinically distinct neurodegenerative diseases by analyzing LRRK2-mediated phosphorylation of Rab12 at amino acid Ser106 (pS106-Rab12). We found that LRRK2 is activated in neurodegenerative diseases and associated with tau pathology, evidenced by elevated Rab12 phosphorylation in DLB subjects with high tau pathology and labeling of GVBs as well as mature tau and α-synuclein pathology in tauopathies and synucleinopathies. For the first time, we provide evidence that LRRK2-mediated hyperphosphorylated Rab proteins represent shared pathological features across synucleinopathies and tauopathies.

## Methods

### Human brain tissue

Brain tissue from 14 neurologically normal control subjects, 5 Alzheimer’s disease (AD) subjects, 17 Dementia with Lewy Bodies (DLB) subjects, 17 Parkinson’s disease (PD) subjects with or without *LRRK2* G2019S mutations, 1 PD subject with a *LRRK2 L1165P* variant, 1 Schizophrenia subject with a *LRRK2 G2019S* mutation, 3 Pick’s disease subjects, 3 Progressive Supranuclear Palsy (PSP) subjects, and 3 Corticobasal Degeneration (CBD) subjects were obtained collectively from the Bryan Brain Bank and Biorepository of the Duke-UNC Alzheimer’s Disease Research Center (ADRC), University of Washington Biorepository and Integrated Neuropathology Laboratory, and the University of Pennsylvania Center for Neurodegenerative Disease Research Brain Bank. Samples included fresh-frozen hippocampus and temporal cortex, as well as slides of formalin-fixed, paraffin embedded (FFPE) brain slices (6µm thickness) of hippocampus (with adjacent entorhinal cortex and temporal cortex), frontal cortex (either inferior frontal cortex or mid frontal gyrus), and/or striatum. All subject information, including tissue samples acquired, proteins analyzed, age, sex, race, cognitive status, post-mortem interval (PMI), *LRRK2* mutation status, and neuropathological assessments can be found in Supplementary Tables 1-4. All donors had provided written informed consent for the use of autopsy material and of clinical and genetic information for research purposes.

### Western blot analysis for human brain tissue

Western blot was performed on hippocampus from 3 control and 7 DLB cases and temporal cortex from 5 control and 11 DLB cases. Brain tissues (∼50mg) were Dounce homogenized in 0.5 mL 1x phosphate-buffered saline (PBS) supplemented with protease inhibitor (Millipore Sigma 4693159001, 1 tablet per 10mL 1x PBS) and phosphatase inhibitor (Millipore Sigma 4906845001, 1 tablet per 10mL 1x PBS) and then centrifuged at 10,000 × g for 10 min at 4°C. Supernatant was discarded, and pellet was Dounce homogenized in 0.4 mL high-salt 1x PBS (500mM NaCl) supplemented with protease and phosphatase inhibitors (both 1 tablet per 10mL 1x PBS), and centrifuged at 10,000 × g for 10 minutes at 4°C. Supernatant was discarded, and pellet was resuspended in 0.4 mL 1% Triton X-100 in 1x PBS supplemented with protease and phosphatase inhibitors (both 1 tablet per 10mL 1x PBS), and centrifuged at 50,000 × g for 20 minutes at 4°C. Saved supernatant was the soluble fraction. Pellet was resuspended in 0.3 mL RIPA (10% SDS, 10% Triton X-100, 10% Sodium deoxycholate in 1x PBS supplemented with protease and phosphatase inhibitors) and sonicated for 10 seconds – resulting in the insoluble fraction. Concentrations of proteins were determined on short-stack SDS-PAGE stain free gels against BSA standards. Proteins were separated by SDS-PAGE and then transferred to PVDF membranes overnight at 25V in tris-glycine transfer buffer with 20% methanol. Membranes were blocked in 5% w/v nonfat dry milk in 1x Tris-buffered saline with 0.1% Tween 20 (TBST) for 30 minutes. Blots were then incubated with primary antibodies (rabbit anti-phospho-Rab12 S106, Abcam ab256487, 1:1000; rabbit anti-Rab12, Proteintech 18843-1-AP, 1:1000; rabbit anti-Hsp70, Thermo Fisher PA5-28003; 1:2000, mouse anti-β-actin, Santa Cruz sc-47778, 1:3000) overnight at 4°C. Phospho-Rab12 antibody was diluted in SignalBoostImmunoreaction Enhancer Kit (Millipore Sigma 407207), and Rab12, Hsp70 and β-actin primary antibodies were diluted in 5% w/v nonfat dry milk in TBST (Tris-buffered saline with 0.1% Tween 20). After washing with 1x TBST, the membrane was incubated in secondary antibody (Peroxidase AffiniPure donkey anti-rabbit, Jackson ImmunoResearch 711-035-152, 1:10,000) for 2 hours at room temperature. Secondary antibody was diluted in SignalBoostImmunoreaction Enhancer Kit for phospho-Rab12 or 5% nonfat dry milk in 1x TBST for Rab12, Hsp70 and β-actin. The signals of target proteins were developed with Crescendo ECL reagent (Millipore) by Chemidoc MP platform (BioRad). Saturated signals on immunoblots were not detected in any experiment used for analysis (ImageLab 6.1). Phospho-Rab12 signal was normalized to Rab12 signal and then quantified relative to control samples. Analysis was performed blinded to group.

### Human brain tissue immunohistochemistry and immunofluorescence

DAB IHC was performed on all cases except one control and one DLB subject (case numbers 38 and 39 in Supplementary Table 1), which were used solely for immunofluorescence (IF). Slides with FFPE brain sections were deparaffinized in xylene (20 minutes), then washed in 100% ethanol (5 minutes), followed by 95% ethanol (5 minutes), and finally distilled H_2_O (10 minutes). Antigen retrieval was then performed by placing slides in boiling citrate buffer (pH 6) for 20 minutes, followed by three H_2_O washes (2 minutes each). Primary antibodies (Rabbit anti-phospho-Rab12 S106, Abcam ab256487, 1:1000; rabbit anti-phospho-LRRK2 S935, Abcam ab133450, 1:1000) were diluted in antibody diluent (Agilent S080983-2) and added to the slides for 45 minutes at 37°C. Slides were washed in H_2_O three times (2 minutes each), and then 1.875% H_2_O_2_ in 100% methanol was added to slides for 8 minutes to block endogenous peroxidase activity. Following three H_2_O washes (2 minutes each), slides were incubated with Dako EnVision^TM^ Dual Link System-HRP (Agilent K400311-2) for 30 minutes at 37°C. Slides were washed with H_2_O three times (2 minutes each) and then incubated with Dako DAB solution (Agilent K346811-2) for 2 minutes. After one H_2_O wash (5 minutes), slides were counterstained with Fisherfinest Hematoxylin+ (ThermoFisher 220-100) for 15-30 seconds, then, following one H_2_O wash (5 minutes), made blue by adding ammonical water (400mL H_2_O with 1mL 28-30% ammonia solution; VWR BDH3014-500MLP) for 15 seconds. Finally, following a 3-minute H_2_O wash, slides were dehydrated through graded alcohols, cleared in xylene, mounted with Permount (Electron Microscopy Sciences 17986-05) and coverslipped. To analyze DAB IHC signal, entire sections were imaged in brightfield on an Olympus SLIDEVIEW VS200 scanning microscope with a 60x (1.42 NA) oil immersion objective. Single-plane images of sections (pixel height and width of 0.0913µm) were exported to QuPath (version 0.5.1), where thresholds for phospho-Rab12- and phospho-LRRK2-positive signal were set and kept consistent across images for each round of IHC. Percent of area positive for phospho-Rab12 and phospho-LRRK2 and density of granulovacuolar degeneration body (GVB)-positive cells labeled by phospho-Rab12 were recorded for each brain region analyzed. Analysis was performed blinded to group.

For IF of 1 control and 1 DLB case (subject numbers 38 and 39 in Supplementary Table 1), slides with FFPE brain sections were deparaffinized in xylene (15 minutes), then washed in 100% ethanol (5 minutes), followed by 95% ethanol (5 minutes), and finally H_2_O (10 minutes). Antigen retrieval was then performed by placing slides in boiling citrate buffer (pH 6) for 20 minutes, followed by an H_2_O wash (10 minutes). Slides were then blocked with 10% normal goat serum in antibody diluent (Agilent S080983-2) for 1 hour, followed by an H_2_O wash (10 minutes). Primary antibodies (Rabbit anti-phospho-Rab12 S106, Abcam ab256487, 1:500; mouse anti-Ck1δ, Santa Cruz sc55553, 1:50; mouse anti-AT8, Thermo Fisher MN1020, 1:500; mouse anti-alpha-synuclein phospho-S129, Abcam ab184674, 1:1000, chicken anti-MAP2, BioLegend 822501, 1:1000) were diluted in antibody diluent (Agilent S080983-2) with 1% normal goat serum and added to the slides for 24 hours at 4°C. Slides were washed in 1x PBS three times (5 minutes each) and then incubated in secondary antibodies (Goat anti-rabbit Alexa Fluor 488, Thermo Fisher A11008, 1:500; goat anti-mouse Alexa Fluor 594, Thermo Fisher A11032, 1:500; goat anti-chicken Alexa Fluor 647, Thermo Fisher A21449, 1:500) diluted in antibody diluent with 1% normal goat serum. Slides were kept in the dark from secondary antibody incubation onward. After three 1x PBS washes (5 minutes each), slides were incubated with TrueBlack (Cell Signaling 92401S) for 30 seconds to reduce tissue autofluorescence. Finally, slides were washed in H_2_O (5 minutes), dried with a kimwipe, mounted with Prolong Diamond Antifade Mountant (Thermo Fisher P36970) and coverslipped (VWR 48393-241). To analyze IF signal, slides were imaged on a Zeiss 780 upright confocal microscope with a 63x (1.4NA) oil immersion objective and 405nm Diode, Argon/2 488nm, 561nm Diode and HeNe 633nm lasers. Images (1024×1024 pixels, pixel height and width of 0.1318µm, 4x pixel averaging) were taken as a z-stack (6 images, 1µm steps) and merged into a maximum intensity projection to create representative images.

### Mice

All animal protocols were approved by Duke University’s Institutional Animal Care and Use Committee. Both male and female mice were employed for all experiments. *Lrrk2* knockout (KO) mice (via deletion of exons 39-40, C57BL/6NJ background) were acquired from Jackson Laboratory (#016121). Both 6-7 month-old wild-type (WT, C57BL/6NJ; n = 4) and *Lrrk2* KO (n = 3) mice were anesthetized with isoflurane, transcardially perfused with 25mL ice-cold normal saline (12.5mL/minute for 2 minutes), then decapitated. Brains were removed, rinsed in ice-cold 1x PBS, and split into individual hemispheres. One hemisphere was immediately microdissected on ice and flash-frozen in liquid nitrogen and stored at -80°C until processed for immunoblotting.

PS19 mice (C57BL/6NJ background), which express P301S mutant human tau under the direction of the mouse prion promoter, *Prnp*, were acquired from Jackson Laboratory (#024841). Young (3-4 months old) and aged (9-10 months old) hemizygous PS19 mice (n = 4 young, 8 aged) and WT littermates (n = 4 young, 5 aged) were anesthetized with isoflurane, transcardially perfused with 25mL ice-cold normal saline (12.5mL/minute for 2 minutes), then decapitated. Brains were removed, rinsed in ice-cold 1x PBS, and split into individual hemispheres. One hemisphere was immediately microdissected on ice to obtain entorhinal cortex. Microdissected brain samples were flash-frozen in liquid nitrogen and stored at -80°C until processed for immunoblotting. The other hemisphere was placed into 4% paraformaldehyde (PFA) in 1x PBS for 24 hours at 4°C, then rinsed with 1x PBS and transferred to 30% sucrose in 1x PBS for at least 72 hours (at 4°C) until slicing.

### Western blot analysis for mouse brain tissue

Cortex from WT (n = 4) and *Lrrk2* KO (n = 3) mice were used for analysis of LRRK2-mediated phosphorylation of Rab12. Entorhinal cortex from WT (n = 4 young, n = 5 aged) and PS19 (n = 4 young, n = 8 aged) mice were used to analyze tau levels in the entorhinal cortex. Microdissected brain samples were weighed and then Dounce homogenized in lysis buffer (94% RIPA buffer, Sigma-Aldrich R0278; 5% protease inhibitor cocktail, Sigma-Aldrich P8340; 1% Halt phosphatase inhibitor cocktail, Thermo Fisher 78420) at a concentration of 10µL buffer per 1mg of brain tissue. After incubating on ice for 10 minutes, homogenized samples were centrifuged at 16,000 × g for 15 minutes at 4°C. Supernatant was collected and stored at -80°C until protein was quantified using a DC protein assay (Bio-Rad 5000112). For western blots, 50µg of protein was loaded for each sample. Protein samples were incubated at 70°C for 10 minutes with NuPAGE sample loading dye (Thermo Fisher NP0007) and dithiothreitol reducing agent (Bio-Rad 1610610). After 4-20% SDS-PAGE (175V, 43 minutes), gels were transferred onto nitrocellulose membranes (Bio-Rad 1704271) using the Trans-Blot Turbo Transfer System (Bio-Rad 1704155) and Trans-Blot Turbo RTA Midi 0.2µm Nitrocellulose Transfer Kit (Bio-Rad 1704271). The blots were blocked in 5% w/v nonfat dry milk in 1x PBST (PBS with 0.05% Tween 20) for 30 minutes. Primary antibodies (Rabbit anti-phospho-Rab12 S106, Abcam ab256487, 1:1000; rabbit anti-Rab12, Proteintech 18843-1-AP, 1:1000; rabbit anti-LRRK2 clone c41-2, Abcam ab133474, 1:2000; mouse anti-AT8, Thermo Fisher MN1020, 1:1000; mouse anti-Tau5, Thermo Fisher AHB0042, 1:5000; mouse anti-GAPDH, Proteintech 60004-1-Ig, 1:5000; rabbit anti-GAPDH, EMD Millipore ABS16, 1:1000) were diluted in 5% nonfat dry milk in 1x PBST, and membranes were incubated in primary antibody for 24 hours at 4°C. Following six 1x PBS washes (5 minutes each), blots were then probed with fluorescent-labeled secondary antibodies (IRDye 680RD donkey anti-mouse, LI-COR 926-68072, 1:10,000; IRDye 680RD donkey anti-rabbit, LI-COR 926-68073, 1:10,000; IRDye 800CW donkey anti-mouse, LI-COR 926-32212, 1:10,000; IRDye 800CW donkey anti-rabbit, LI-COR 926-32213, 1:10,000), diluted in 5% nonfat dry milk in 1x PBST, for 45 minutes (kept in the dark from secondary antibody incubation through imaging). After three 1x PBS washes (5 minutes each), blots were scanned using an Odyssey Imaging scanner (LI-COR). Fluorescence intensities were quantified using ImageStudio Lite Software (LI-COR), and the signal from proteins of interest were normalized to GAPDH. To quantify Rab12 phosphorylation, phospho-Rab12 signal and Rab12 signal were both normalized to their respective GAPDH signals. Then, normalized phospho-Rab12 was divided by normalized Rab12 to give a phospho-Rab12/Rab12 ratio. Analysis was performed blinded to group.

### Immunofluorescence of mouse brain tissue

Brains from WT (n = 5 young, 6 aged) and PS19 (n = 6 young, 9 aged) mice were sliced on a Leica microtome (40µm thickness), and slices were stored in 50% glycerol/50% 1x PBS at -20°C until IF was performed. For each animal, free-floating brain slices (2 slices per region analyzed for each experiment) were washed in 1x PBS two times (5 minutes each) and then transferred into blocking solution (10% normal donkey serum in 1x PBS with 0.3% Triton X-100) for 1 hour. After three 1x PBS washes (5 minutes each), slices were incubated with TrueBlack (Cell Signaling 92401S) for 30 seconds to reduce tissue autofluorescence. Following two 5-minute 1x PBS washes, slices were then incubated in primary antibodies (Rabbit anti-phospho-Rab12 S106, Abcam ab256487, 1:500; mouse anti-Ck1δ, Santa Cruz sc55553, 1:50; mouse anti-AT8, Thermo Fisher MN1020, 1:500; rat anti-LAMP1, Developmental Studies Hybridoma Bank 1D4B, 1:250; rabbit anti-phospho-ubiquitin S65, Sigma-Aldrich ABS1513-I 1:1000; chicken anti-NeuN, GeneTex 00837, 1:1000) diluted in 1% normal donkey serum in 1x PBS for 24 hours at 4°C. Slices were washed in 1x PBS three times (10 minutes each) and then incubated in secondary antibodies (Donkey anti-rabbit Alexa Fluor 488, Thermo Fisher A21206, 1:500; donkey anti-mouse Alexa Fluor 594, Thermo Fisher A21203, 1:500; donkey anti-rat Alexa Fluor 594, Thermo Fisher A21209, 1:500; donkey anti-mouse Alexa Fluor 647, Thermo Fisher A31571, 1:500; donkey anti-chicken Alexa Fluor 647, Thermo Fisher A78952, 1:500) diluted in 1% normal donkey serum in 1x PBS for 1 hour. Slices were kept in the dark from secondary antibody incubation onward. Following a 10-minute 1x PBS wash, slices were incubated with Hoechst 33342 (1:5000 in 1x PBS, Thermo Fisher H3570) for 5 minutes. Finally, slices were washed in 1x PBS (10 minutes), transferred onto slides, dried with a Kimwipe, mounted with Prolong Diamond Antifade Mountant (Thermo Fisher P36970) and coverslipped. To assess the phospho-state-specific nature of labeling (as performed in [11]), after the initial 1x PBS wash, some slices were incubated for 10 minutes in 1x NEBuffer with 1mM MnCl_2_ provided by the Lambda Protein Phosphatase kit (New England Biolabs P0753S). Slices were then incubated in 1x NEBuffer with 1mM MnCl_2_, either with or without 10,000 units/mL Lambda phosphatase, for 6 hours at 37°C. Slices were then washed in 1x PBS and then transferred into blocking solution, and IF was performed as described above.

To analyze IF signal, slides were imaged on a Zeiss 780 upright confocal microscope with a 63x (1.4NA) oil immersion objective and 405nm Diode, Argon/2 488nm, 561nm Diode and HeNe 633nm lasers. For quantification, images (1024×1024 pixels, pixel height and width of 0.1318µm, 4x pixel averaging) were taken as a z-stack (6 images, 1µm steps). To display granulovacuolar degeneration body ultrastructure and co-localization of markers in granulovacuolar degeneration bodies, some high-resolution images (2048×2048 pixels, pixel height and width of 0.0659µm, 4x averaging) were also taken as a z-stack (6 images, 1µm steps). Both single-plane images and average intensity projections of z-stacks are shown as representative images. For analysis, 3 z-stack images were taken for each hippocampus slice (6 total images per animal) and 5 z-stack images were taken for each entorhinal cortex slice (10 total images per animal). All images were imported into QuPath (version 0.5.1) for quantification. A threshold for AT8-positive signal was set and kept consistent across images, and AT8 integrated density was calculated by multiplying AT8-positive area and mean pixel intensity of AT8-positive signal. For quantification of phospho-Rab12 puncta in AT8-positive neurons, “Low AT8” neurons were those displaying puncta-like labeling of AT8 (Fig. 5a), while “High AT8” neurons were those displaying dense AT8 positive labeling throughout the cytoplasm of the neuron (i.e., “pre-tangle” levels of AT8, Fig. 5b). For all other mouse IF analyses, the presence of at least 1 Ck1δ puncta in a NeuN-positive cell was deemed a GVB-positive neuron. Phospho-Rab12, Ck1δ, and phospho-ubiquitin puncta were counted in neurons with Ck1δ puncta and co-localization of these puncta were manually recorded. phospho-Rab12 and Ck1δ puncta were also counted for localization with LAMP1 in Ck1δ-positive neurons, with the presence of a LAMP1-positive structure surrounding phospho-Rab12 and/or Ck1δ puncta being deemed co-localization of LAMP1 with those structures.

### Statistical analyses

Data were analyzed in GraphPad Prism (version 10.0.2). Human subject characteristics (age, PMI, and neuropathological stages) were compared across diagnoses via a one-way analysis of variance (ANOVA). Non-parametric Spearman tests of correlations and Kruskal-Wallis H-tests with Dunn’s test for multiple comparisons were used for human experiments, as most human measures were not normally distributed. Mouse studies were analyzed via Student’s two-tailed unpaired *t* test or two-way ANOVA with Bonferroni’s post hoc multiple comparisons test. Significance was defined by p-value < 0.05. For all graphs, bars represent mean ± standard error of the mean (SEM). All data points presented in graphs represent the analysis of an independent sample or subject.

## Results

### Aberrant Rab12 phosphorylation in DLB

We sought to determine whether Rab12 phosphorylation mediated by LRRK2 is altered in neurodegenerative diseases with tau and/or α-synuclein pathology. To replicate previous findings that LRRK2 is the primary kinase for Rab12 phosphorylation in the brain, the cortex was dissected from wild-type (WT) and *Lrrk2* knockout (KO) mice and subjected to quantitative immunoblotting for pS106-Rab12 and Rab12. Consistent with previous reports, we observed in cortex derived from *Lrrk2* KO mice a significant decrease in pS106-Rab12 compared to WT (Supplementary Fig. 1a, b) [47, 51, 53, 80, 81, 91].

We first focused on DLB, which despite being characterized by the presence of Lewy bodies, often displays moderate to severe 3R/4R tau pathology post-mortem and therefore is a relevant disease to interrogate α-synuclein and tau co-pathologies [13, 33, 37, 76]. The hippocampus and temporal cortex were chosen for analysis because they display high levels of tau pathology and manifest tau pathology early in disease progression, both in DLB and AD [9, 10, 16, 90]. Brains from subjects with DLB with a range of AD co-pathology levels, and unaffected controls, were subjected to quantitative immunoblotting for pS106-Rab12 and Rab12 (Supplementary Table 1). Positive and specific pS106-Rab12 signal was observed in the soluble fraction (Fig. 1a, b), but not in the insoluble fraction, where only non-specific signal was observed (Supplementary Fig. 2a). We discovered that pS106-Rab12 was ∼7-fold higher in hippocampus derived from subjects with DLB, when stratified by Braak neurofibrillary tangle stage (Braak > 3), relative to controls (Fig. 1a, c). Rab12 phosphorylation was also elevated by >20-fold in temporal cortex of DLB cases (Braak > 3) when compared to either control or DLB cases (Braak ≤ 3) (Fig. 1b, d).

**Fig. 1.**
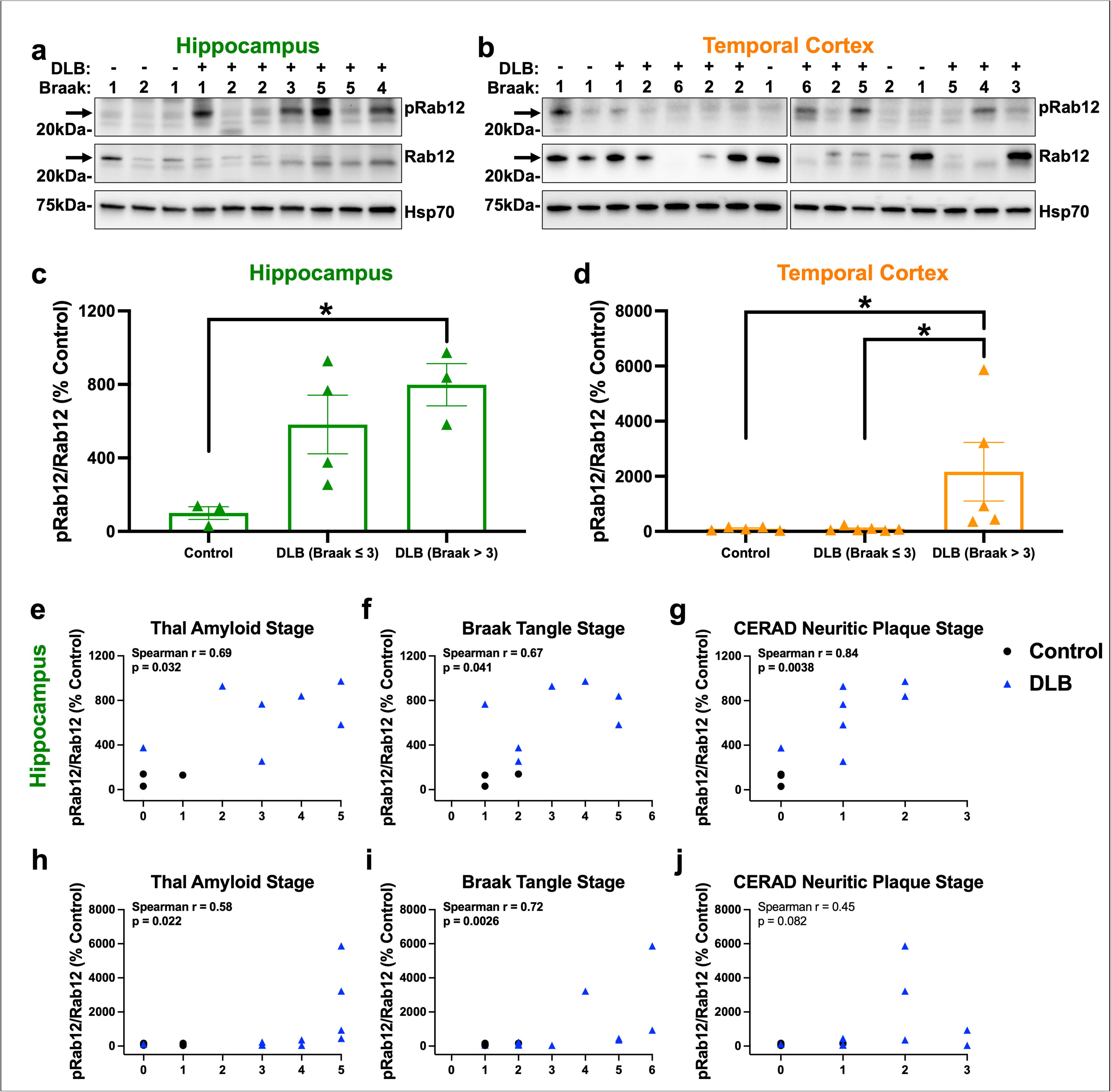
pS106-Rab12 quantitative immunoblotting in DLB hippocampus and temporal cortex. Representative western blot of (**a**) hippocampus derived from control (n = 3) and DLB (n = 7) subjects or (**b**) temporal cortex derived from control (n = 5) and DLB (n = 11) subjects, assessed for pS106-Rab12 (pRab12), Rab12, and Hsp70 as a loading control. Braak tangle stage (1-6) of each control or DLB subject is shown. (**c, d**) Quantification of western blots show that pRab12 is elevated in both the hippocampus and temporal cortex in DLB (Braak > 3) cases compared to controls. pRab12 is also elevated in temporal cortex in DLB (Braak > 3) cases compared to DLB (Braak ≤ 3) cases. Data is presented as pRab12/Rab12 (% control). (**e-j**) Correlations (with Spearman correlation coefficients and associated p-values) of (**e-g**) hippocampus and (**h-j**) temporal cortex pRab12/Rab12 with (**e, h**) Thal amyloid stage, (**f, i**) Braak tangle stage, and (**g, j**) CERAD neuritic plaque stage in control and DLB subjects (significant correlations are bolded). Data are presented as mean ± SEM, and each point represents an individual subject. *p < 0.05 via Kruskal-Wallis H-test with Dunn’s post hoc multiple comparisons test

Consistent with a relationship between Rab12 phosphorylation and tau pathology, we found that pS106-Rab12 phosphorylation was positively correlated with AD co-pathology measures (Fig. 1e-j). pS106-Rab12 was positively and moderately correlated with Thal amyloid and Braak neurofibrillary tangle (Braak tangle) stages, and strongly correlated with CERAD neuritic plaque pathology stage, in the hippocampus of control and DLB cases (Fig. 1e-g). In the temporal cortex, pS106-Rab12 was positively and moderately correlated with Thal amyloid and Braak tangle stages, but not with CERAD neuritic plaque stage, in control and DLB cases (Fig. 1h-j). Rab12 phosphorylation was not significantly correlated with age or post-mortem interval (PMI) in either the hippocampus or temporal cortex (Supplementary Fig. 2b-e). These findings demonstrate that pS106-Rab12, which is an activator of LRRK2 and proxy for LRRK2 kinase activity, is elevated in DLB cases with more advanced amyloid-β and tau pathology.

### pS106-Rab12 localizes to granulovacuolar degeneration bodies in DLB and AD

To further investigate the elevated levels of phosphorylated Rab12 described in DLB (Fig. 1), we next determined the localization of pS106-Rab12 by IHC in the hippocampus, adjacent temporal cortex (comprising both entorhinal cortex and temporal cortex) and frontal cortex from DLB and control cases (subject characteristics can be found in Supplementary Table 1). There was no significant difference in age or PMI between groups (Supplementary Table 2). We observed pS106-Rab12 labeling features highly reminiscent of granulovacuolar degeneration bodies (GVBs) across subjects (Fig. 2a), with significantly higher pS106-Rab12 GVB-like cell density in subjects with DLB when stratified by Braak neurofibrillary tangle stage (Braak > 3), relative to DLB (Braak ≤ 3) or control cases (Fig. 2a, b).

**Fig. 2.**
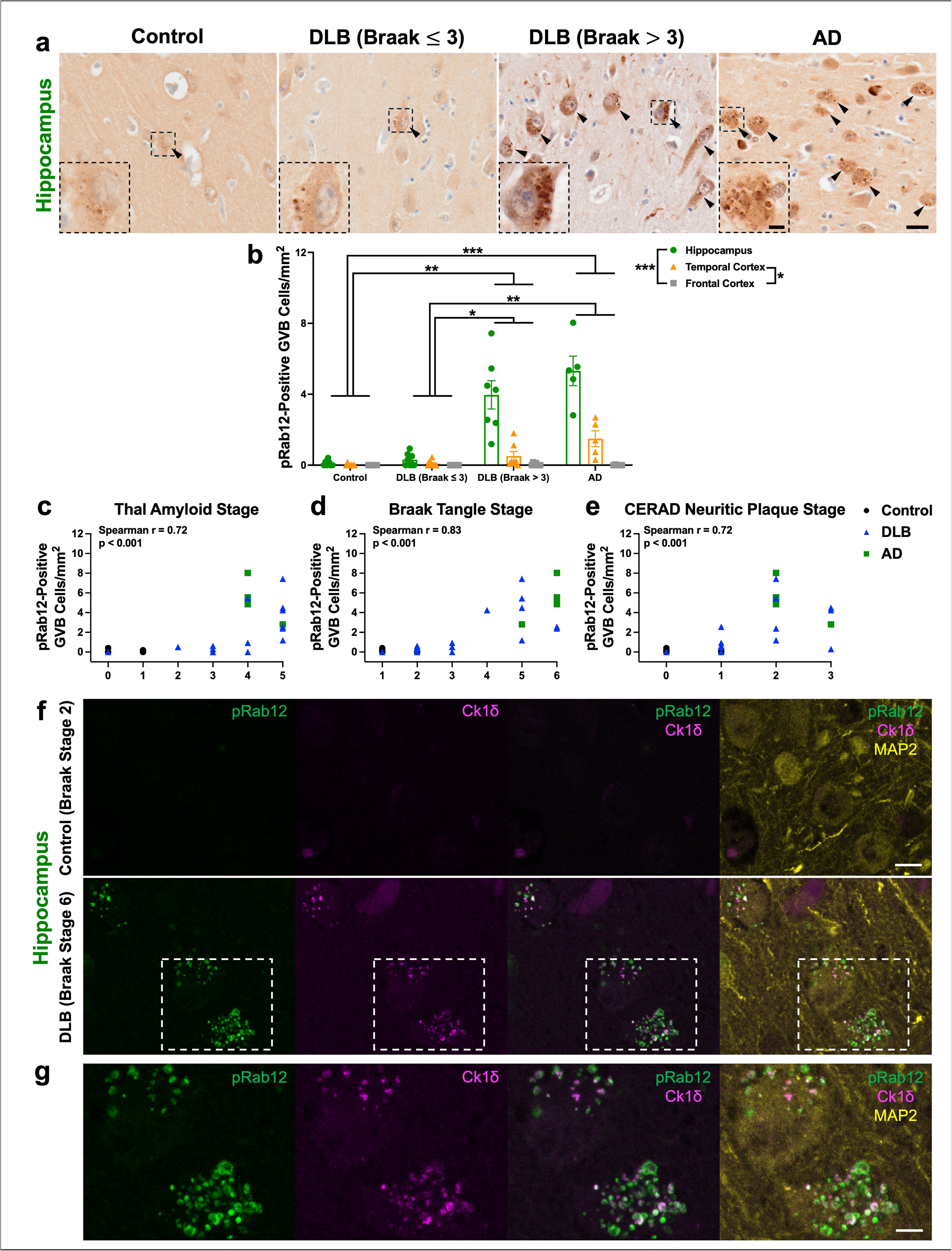
pS106-Rab12 IHC in DLB and AD brains reveals pS106-Rab12 labeling of GVBs. (**a**) Representative 60x DAB IHC images and insets of the hippocampus from a control, DLB (Braak ≤ 3), DLB (Braak > 3), and AD case. Tissue was incubated with an antibody against pS106-Rab12. Black arrows denote pS106-Rab12 GVB-positive cells. (**b**) Quantification of the pS106-Rab12-positive GVB cell density in the hippocampus, temporal cortex, and frontal cortex of control (n = 7), DLB (Braak ≤ 3, n = 9), DLB (Braak > 3, n = 7), and AD (n = 5) cases. (**c-e**) Correlations (with Spearman correlation coefficients and associated p-values) of hippocampus pS106-Rab12 GVB-labeled cell density with (**c**) Thal amyloid stage, (**d**) Braak tangle stage, and (**e**) CERAD neuritic plaque stage in control, DLB and AD cases (significant correlations are bolded). (**f**) Representative 63x IF images of the hippocampus from a control (Braak tangle stage 2) and a DLB (Braak tangle stage 6) case. Tissue was incubated against pS106-Rab12 (pRab12, green), Ck1δ (magenta), and neuronal marker MAP2 (yellow). (**g**) Inset of (**f**) showing co-localization of pRab12 and Ck1δ in hippocampus tissue from a DLB subject. Data are presented as mean ± SEM and each point represents an individual subject. *p < 0.05, **p < 0.01, ***p < 0.001 via Kruskal-Wallis H-test with Dunn’s post hoc multiple comparisons test. Scale bar in (**a**) = 20µm, inset scale bar in (**a**) = 5µm, scale bar in (**f**) = 10µm, scale bar in (**g**) = 5µm

To determine if this pS106-Rab12 labeling is specific to DLB or common across neurodegenerative diseases, we also analyzed hippocampus, temporal cortex and frontal cortex derived from AD subjects. All AD cases had Braak neurofibrillary tangle pathology stages of 5 or above (shown by labeling of phospho-tau S202/T205 by AT8, Supplementary Fig. 3). We found that like DLB, pS106-Rab12 GVB-like cell density was significantly higher in AD cases compared to DLB (Braak ≤ 3) and control cases (Fig. 2a, b). Interestingly, we observed regional differences in pS106-Rab12 GVB-pattern labeling. The highest density of GVB-like labeling was observed in the hippocampus, with the lowest density in the frontal cortex (Fig. 2a, b, Supplementary Fig. 4a). This pattern recapitulates regional differences in GVB density and tau pathology in AD and DLB [9, 10, 16, 82, 90]. Consistent with the link between GVBs and tau pathology, we observed a significant and positive correlation of pS106-Rab12 GVB-like cell density with Thal amyloid, Braak tangle, and CERAD neuritic plaque stages in the hippocampus of DLB and AD cases (Fig. 2c-e). Hippocampus pS106-Rab12 GVB-like cell density was not significantly correlated with age or PMI (Supplementary Fig. 4b, c).

Co-localization analysis demonstrates that pS106-Rab12 overlaps with the canonical GVB-specific marker casein kinase 1δ (Ck1δ, Fig. 2f, g) [94]. Of note, we found Ck1δ- and pS106-Rab12-positive GVBs only in the DLB case, but not the control, consistent with minimal GVB formation in the absence of disease (Fig. 2f, g) [94]. Though the majority of pS106-Rab12 and Ck1δ signal showed strong overlap, pS106-Rab12-positive/Ck1δ-negative and pS106-Rab12-negative/Ck1δ-positive puncta were observed (Fig. 2f, g). In all, these findings show that pS106-Rab12 preferentially labels GVBs that accumulate in neurodegenerative diseases with tau pathology.

### pS106-Rab12 labels GVBs in both genetic and idiopathic PD

We next performed IHC for pS106-Rab12 in both the hippocampus and adjacent temporal cortex of a cohort of PD cases that harbor the G2019S pathogenic mutation (LRRK2^GS^ PD), compared to idiopathic PD (iPD) and unaffected controls. All iPD and LRRK2^GS^ PD cases were diagnosed with PD, while 6 out of 10 iPD and 2 out of 7 LRRK2^GS^ PD subjects had PD dementia (PDD) (Supplementary Table 3). AT8 labeling was used for Braak neurofibrillary tangle staging and stratifying LRRK2^GS^ PD and iPD cases into (Braak ≤ 3) and (Braak > 3) groups (Supplementary Fig. 5, all subject characteristics can be found in Supplementary Table 3). There was no significant difference in age or PMI between groups (Supplementary Table 4).

We observed pS106-Rab12 labeling of GVBs in the hippocampus and temporal cortex across subjects (Fig. 3a, Supplementary Fig. 6). Significantly elevated pS106-Rab12 GVB cell density was detected in LRRK2^GS^ PD cases when stratified by Braak neurofibrillary tangle stage (Braak > 3) compared to controls (Fig. 3a, b). pS106-Rab12 GVB cell density was increased to a similar magnitude in iPD when stratified by Braak neurofibrillary tangle stage (Braak > 3) relative to controls (Fig. 3a, b). pS106-Rab12 GVB cell density in iPD (Braak > 3) cases was significantly elevated compared to controls when only hippocampus was analyzed (Fig. 3b), though the comparison between iPD (Braak > 3) and controls did not reach statistical significance when all brain regions were considered. Consistent with GVB staging in AD, pS106-Rab12 GVB labeling was less elevated in the temporal cortex in LRRK2^GS^ PD and iPD compared to the hippocampus (Fig. 3a, b, Supplementary Fig. 6) [82]. Importantly, we observed a significant positive correlation of hippocampus pS106-Rab12 GVB cell density with Braak tangle stage, but not Thal amyloid stage or CERAD neuritic plaque stage across control, LRRK2^GS^ PD and iPD cases (Fig. 3c-e). A positive correlation of pS106-Rab12 GVB cell density with age was also observed, although this correlation was much weaker than the correlation with Braak tangle stage (Supplementary Fig. 7a). There was no significant correlation of hippocampus pS106-Rab12 GVB cell density with PMI (Supplementary Fig. 7b).

**Fig. 3.**
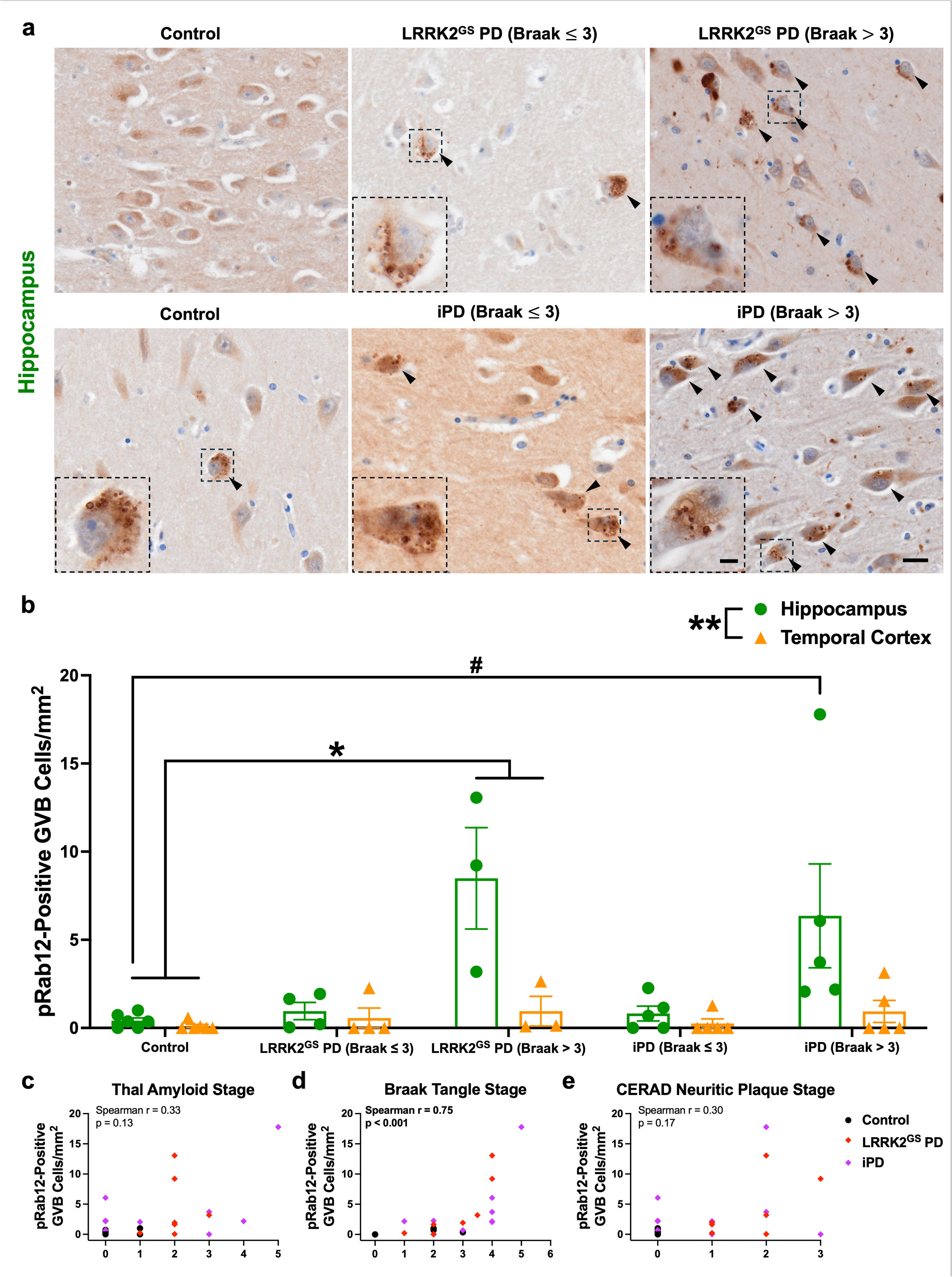
pS106-Rab12 GVB labeling in LRRK2^GS^ PD and iPD. (**a**) Representative 60x images and insets of the hippocampus from a control, LRRK2^GS^ PD (Braak ≤ 3), LRRK2^GS^ PD (Braak > 3), iPD (Braak ≤ 3), and iPD (Braak > 3) case. Tissue was incubated with an antibody against pS106-Rab12. Black arrows denote pS106-Rab12 GVB-positive cells. (**b**) Quantification of the pS106-Rab12 GVB-labeled cell density in the hippocampus and temporal cortex of control (n = 6), LRRK2^GS^ (Braak ≤ 3, n = 4), LRRK2^GS^ (Braak > 3, n = 3), iPD (Braak ≤ 3, n = 5), and iPD (Braak > 3, n = 5) cases. (**c-e**) Correlations (with Spearman correlation coefficients and associated p-values) of hippocampus pS106-Rab12 GVB-labeled cell density with (**c**) Thal amyloid stage, (**d**) Braak tangle stage, and (**e**) CERAD neuritic plaque stage in control, LRRK2^GS^ PD, and iPD cases (significant correlations are bolded). Data are presented as mean ± SEM and each point represents an individual subject. *p < 0.05 compared to controls, **p < 0.01 between regions, ^#^p < 0.05 for hippocampus compared to control hippocampus via Kruskal-Wallis H-test with Dunn’s post hoc multiple comparisons test. Scale bar in (**a**) = 20µm, inset scale bar = 5µm

We also explored the *LRRK2* L1165P variant which is exceedingly rare and detected in a single early-onset PD case with unclear causality, as well as a LRRK2^GS^ subject with Schizophrenia and no PD diagnosis which was reported to have no pathology (α-synuclein or tau) [17, 32]. The LRRK2^L1165P^ PD subject and LRRK2^GS^ Schizophrenia subject both had Braak neurofibrillary tangle stages of 1-2 (Supplementary Table 3). We found pS106-Rab12 GVB labeling in the LRRK2^L1165^ PD case, but not the LRRK2^GS^ Schizophrenia case (Supplementary Fig. 8).

Taken together, these findings show that, similar to DLB and AD, pS106-Rab12 co-localizes with GVBs in LRRK2^GS^ PD and iPD, and phosphorylated Rab12 GVB cell density is correlated with levels of tau pathology. This suggests that pS106-Rab12 labeling of GVBs is consistently observed across neurodegenerative diseases with tau pathology, regardless of *LRRK2* mutation status.

### pS106-Rab12 GVBs accumulate with age in the PS19 tauopathy mouse model

To extend our human brain post-mortem findings to a mouse model with predictable pathological progression of tau, we used the well-studied PS19 strain of mice, which overexpress the P301S pathogenic mutant human tau in neurons and recapitulate hallmarks of tauopathy, including progressive tau pathology, cognitive dysfunction, and neurodegeneration [84, 96]. Consistent with previous studies, the PS19 mice display tau hyperphosphorylation as observed in tauopathies, evidenced by increased IF of AT8 in hippocampus and entorhinal cortex in aged PS19 mice compared to WT (Fig. 4a, b). AT8 IF signal was significantly higher in the entorhinal cortex compared to the hippocampus of PS19 mice (Fig. 4b). Further, we performed quantitative western blotting and observed increased AT8, at molecular weights representing human hyperphosphorylated tau, in entorhinal cortex derived from both young (3-4 month-old) and aged (9-10 month-old) PS19 mice compared to WT (Supplementary Fig. 9a, b). The increase in hyperphosphorylated tau was associated with a concomitant increase in total tau detected by the antibody Tau5 in both young and aged PS19 relative to WT mice (Supplementary Fig. 9c).

**Fig. 4.**
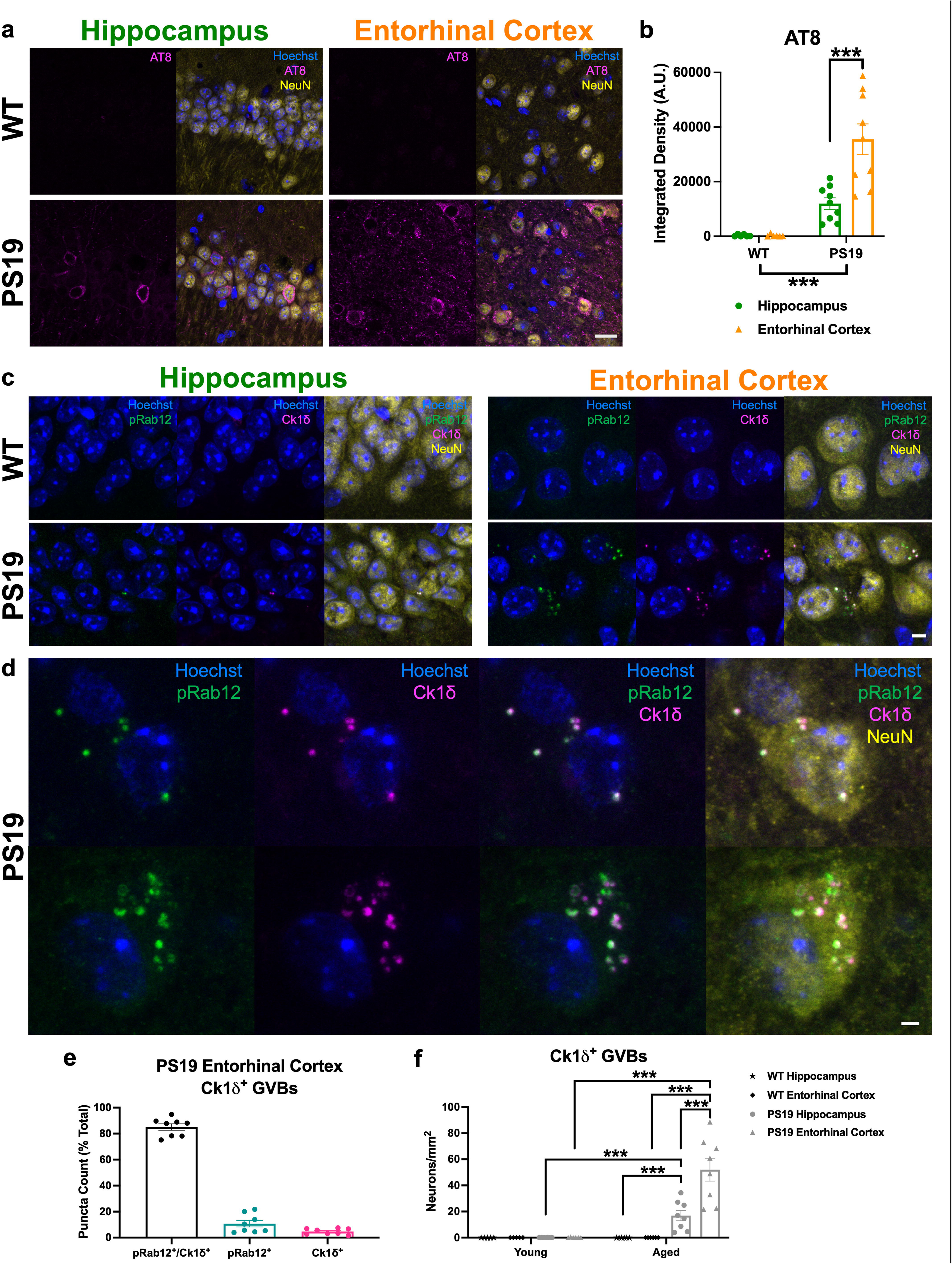
pS106-Rab12 labels GVBs in aged PS19 mouse brain. (**a**) Representative 63x images of Hoechst (blue), AT8 (magenta) and NeuN (yellow) in hippocampus and entorhinal cortex of aged (9-10 month-old) WT and PS19 mice. (**b**) Quantification of AT8 integrated density in hippocampus and entorhinal cortex of aged WT (n = 6) and PS19 (n = 9) mice. (**c**) Representative 63x images of Hoechst (blue), pS106-Rab12 (pRab12, green), Ck1δ (magenta), and neuronal marker NeuN (yellow) in hippocampus and entorhinal cortex of aged WT and PS19 mice. (**d**) Zoomed-in 63x representative images of aged PS19 entorhinal cortex showing overlap of pRab12 and Ck1δ puncta. (**e**) Quantification of pRab12 and Ck1δ puncta co-localization in aged PS19 entorhinal cortex (n = 8). (**f**) Quantification of Ck1δ GVB neuron density in young (3-4 month-old) and aged WT (n = 5 young, 6 aged) and PS19 (n = 6 young, 8 aged) hippocampus and entorhinal cortex. Data are presented as mean ± SEM and each point represents an individual animal. ***p < 0.001 via two-way ANOVA with Bonferroni’s post hoc multiple comparisons test. Scale bar in (**a**) = 20µm, scale bar in (**c**) = 5µm, scale bar in (**d**) = 2µm

In hippocampus and entorhinal cortex derived from aged PS19 mice, we observed neuronal, pS106-Rab12-positive GVB labeling, shown by overlap with GVB marker Ck1δ, that was not observed in WT mice (Fig. 4c, d). Quantification of pS106-Rab12- and Ck1δ-labeled puncta in these neurons demonstrated high co-localization, with 85% of puncta in GVB-labeled neurons displaying labeling of both pS106-Rab12 and Ck1δ (Fig. 4e). Significantly more GVB-labeled neurons were observed in aged PS19 compared to aged WT mice, in which Ck1δ labeling was not detectable (Fig. 4f). These pS106-Rab12 GVBs accumulated with age, as GVB-positive neuron density was significantly higher in aged PS19 compared to young PS19 mice (Fig. 4f, Supplementary Fig. 9d). Similar to regional differences in AT8 signal, greater GVB neuron density was observed in the entorhinal cortex compared to hippocampus in aged PS19 mice (Fig. 4f), suggesting a correlation between hyperphosphorylated tau accumulation and GVB-labeled neuron density. Notably, while young PS19 mice displayed almost no detectable GVB labeling (Fig. 4f, Supplementary Fig. 9d), the rare GVB-positive neurons observed displayed pS106-Rab12-positive GVBs (Supplementary Fig. 9e).

Consistent with previously published work showing pS106-Rab12 IF detects a phospho-epitope, phosphatase (PPase) treatment in PS19 brain slices before staining abolished pS106-Rab12 signal, while preserving the Ck1δ signal, which does not detect a phospho-epitope (Supplementary Fig. 10a, b) [11]. PPase treatment similarly abolished the signal of AT8, which detects S202/T205 phospho-epitopes of tau (Supplementary Fig. 11a, b). Together, these findings demonstrate 1) the majority of GVBs are pS106-Rab12-positive, and 2) pathological tau initiates accumulation of phosphorylated Rab12-positive GVBs in an age-dependent manner.

### Pathological tau accumulation-initiated pS106-Rab12-positive GVBs are lysosomal structures

We next sought to investigate the relationship between hyperphosphorylated tau levels and pS106-Rab12 GVB labeling within each neuron in PS19 mice. We found that pS106-Rab12 puncta co-localized with AT8 in aged PS19 entorhinal cortex neurons where AT8 levels were low and displayed puncta-like labeling of AT8 (Fig. 5a). We also discovered that the number of pS106-Rab12 puncta found in these pS106-Rab12-positive neurons is significantly higher in neurons with high AT8 labeling (i.e., “pre-tangle” tau pathology) compared to neurons with comparatively low AT8 labeling (Fig. 5a-c).

**Fig. 5.**
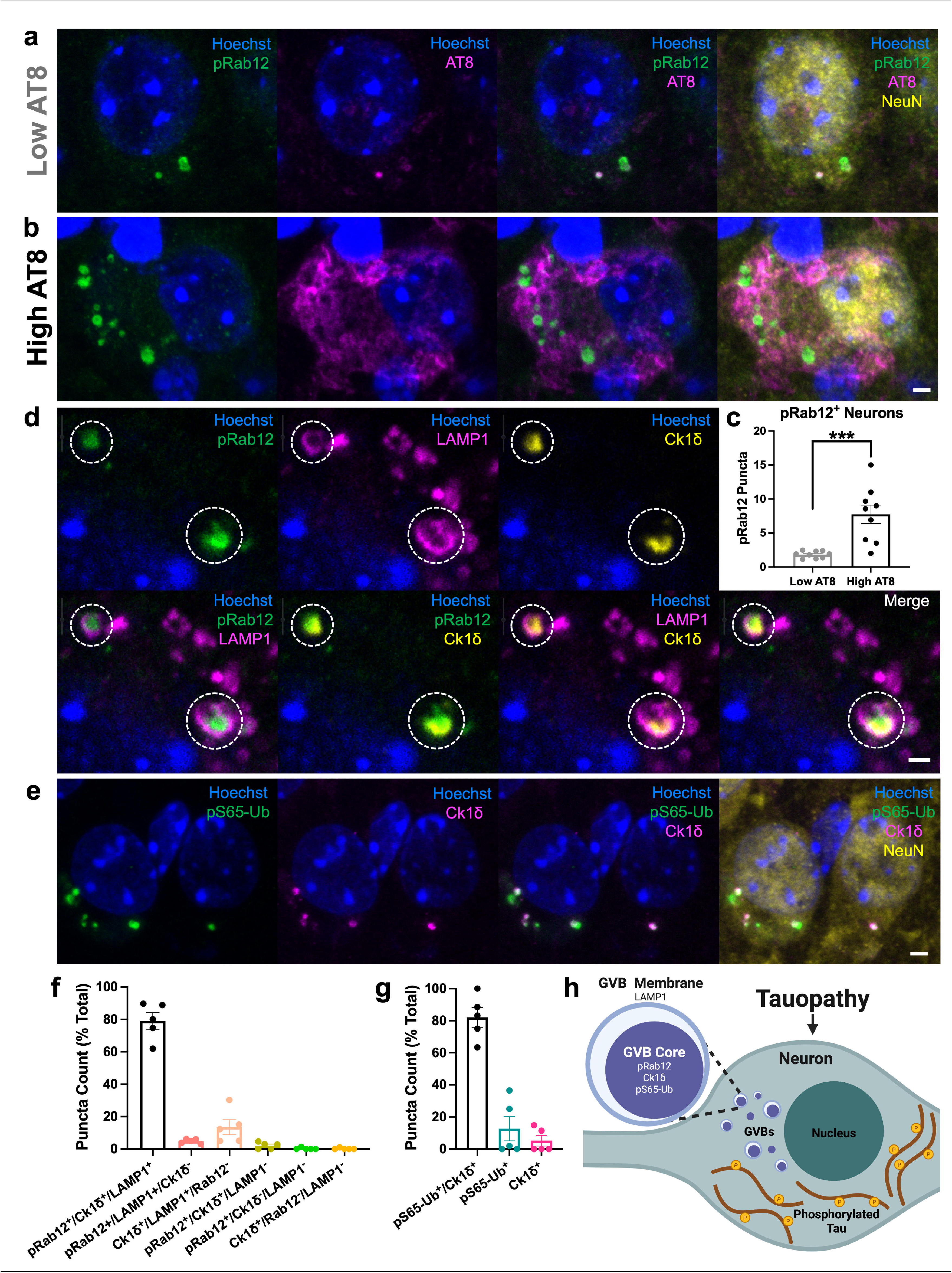
pS106-Rab12-positive GVBs are lysosomal structures that also contain mitophagy protein markers. (**a, b**) Representative 63x images of Hoechst (blue), pS106-Rab12 (pRab12, green), AT8 (magenta) and NeuN (yellow) in aged PS19 entorhinal cortex, showing pRab12 puncta in neurons with (**a**) low and (**b**) high AT8 signal. (**c**) Quantification of pRab12 puncta number in aged PS19 mouse (n = 9) entorhinal cortex neurons with low or high AT8 signal. (**d**) Representative 63x single-plane images of Hoechst (blue), pRab12 (green), LAMP1 (magenta) and Ck1δ (yellow) in aged PS19 entorhinal cortex. Dashed circles show pRab12- and Ck1δ-positive dense GVB cores surrounded by a LAMP1-positive outer membrane. (**e**) Representative 63x images of Hoechst (blue), pS65-Ub (green), Ck1δ (magenta), and NeuN (yellow) in aged PS19 entorhinal cortex showing overlap of pS65-Ub and Ck1δ puncta. (**f**) Quantification of pRab12, Ck1δ, and LAMP1 co-localization in aged PS19 entorhinal cortex (n = 5) showing the majority of pRab12 and Ck1δ puncta in neurons with GVBs are triple-positive for pRab12, Ck1δ and LAMP1. (**g**) Quantification of pS65-Ub and Ck1δ co-localization in aged PS19 entorhinal cortex (n = 5). (**h**) Schematic showing that in tauopathies, GVBs form in neurons with early hyperphosphorylated tau accumulation, and the dense core of these GVBs contain pRab12. Data are presented as mean ± SEM, and each point represents an individual animal. ***p < 0.001 via unpaired Student’s *t* test. Scale bar in (**a**) = 2µm, scale bar in (**d**) = 1µm, scale bar in (**e**) = 2µm. Schematic created with BioRender.com

Previous work has demonstrated that GVBs are lysosomal structures with a dense core (labeled by Ck1δ) surrounded by an outer membrane that is labeled by lysosomal membrane markers like lysosomal-associated membrane protein 1 (LAMP1) [42, 95]. Therefore, we performed IF co-labeling of LAMP1 with pS106-Rab12 and Ck1δ in aged PS19 mouse entorhinal cortex. We found that, consistent with GVBs, Ck1δ labeled the GVB dense core, which was surrounded by a LAMP1-positive membrane (Fig. 5d, f, Supplementary Fig. 12). pS106-Rab12, consistent with its co-labeling of Ck1δ, labeled the dense core of GVBs (Fig. 5d, Supplementary Fig. 12). Consistent with quantification described in (Fig. 4e), 79% of observed pS106-Rab12 and Ck1δ puncta in neurons with Ck1δ-positive GVBs were triple positive for pS106-Rab12, Ck1δ, and LAMP1 (Fig. 5f).

In addition to displaying lysosomal markers, GVBs have been previously reported to co-localize with the mitophagy marker phospho-Ubiquitin S65 (pS65-Ub), and pathogenic *LRRK2* mutations disrupt mitophagy [35, 36, 65, 75]. We observed co-labeling of pS65-Ub and Ck1δ, replicating this finding (Fig. 5e, g). 82% of observed puncta in GVB-positive neurons were double-positive for pS65-Ub and Ck1δ (Fig. 5g). Collectively, these findings demonstrate that pS106-Rab12-labeled GVBs are lysosomal structures that also contain mitophagy-associated markers. Overall based on these mouse and human findings, we demonstrate pS106-Rab12 labeling of GVBs in neurons, linking LRRK2 kinase activation, lysosomes and tau pathology (Fig. 5h).

### pS106-Rab12 labels pathological protein inclusions across neurodegenerative diseases

To determine whether pS106-Rab12 associates with mature protein pathology in neurodegenerative diseases, pS106-Rab12 labeling of mature tau and α-synuclein inclusions was analyzed. In the DLB and AD cohort, pS106-Rab12 labeled pathological tau inclusions, in the form of neurofibrillary tangles and dystrophic neurites, in both the hippocampus and temporal cortex of DLB (Braak > 3) and AD cases (Fig. 6a). pS106-Rab12 labeled occasional dystrophic neurites in the frontal cortex of DLB (Braak > 3) and AD cases as well (Supplementary Fig. 13a). pS106-Rab12 also appeared to label Lewy bodies in DLB, independent of Braak stage (Fig. 6a, Supplementary Fig. 13a). Group comparisons revealed significantly higher pS106-Rab12-positive area in the hippocampus, entorhinal cortex and frontal cortex of DLB and AD cases compared to controls (Fig. 6b). Consistent with labeling of tau pathology, there was a regional difference in pS106-Rab12-positive area across all cases analyzed, with significantly higher positive area in the hippocampus compared to the frontal cortex (Fig. 6b). We found a significant positive correlation of hippocampus pS106-Rab12-positive area with Thal amyloid, Braak neurofibrillary tangle, and CERAD neuritic plaque pathology stages (Fig. 6c-e). Notably, the strongest correlation was with Braak neurofibrillary tangle stage (Fig. 6c-e). There was no significant correlation of hippocampus pS106-Rab12-positive area with age or PMI (Supplementary Fig. 13b, c).

**Fig. 6.**
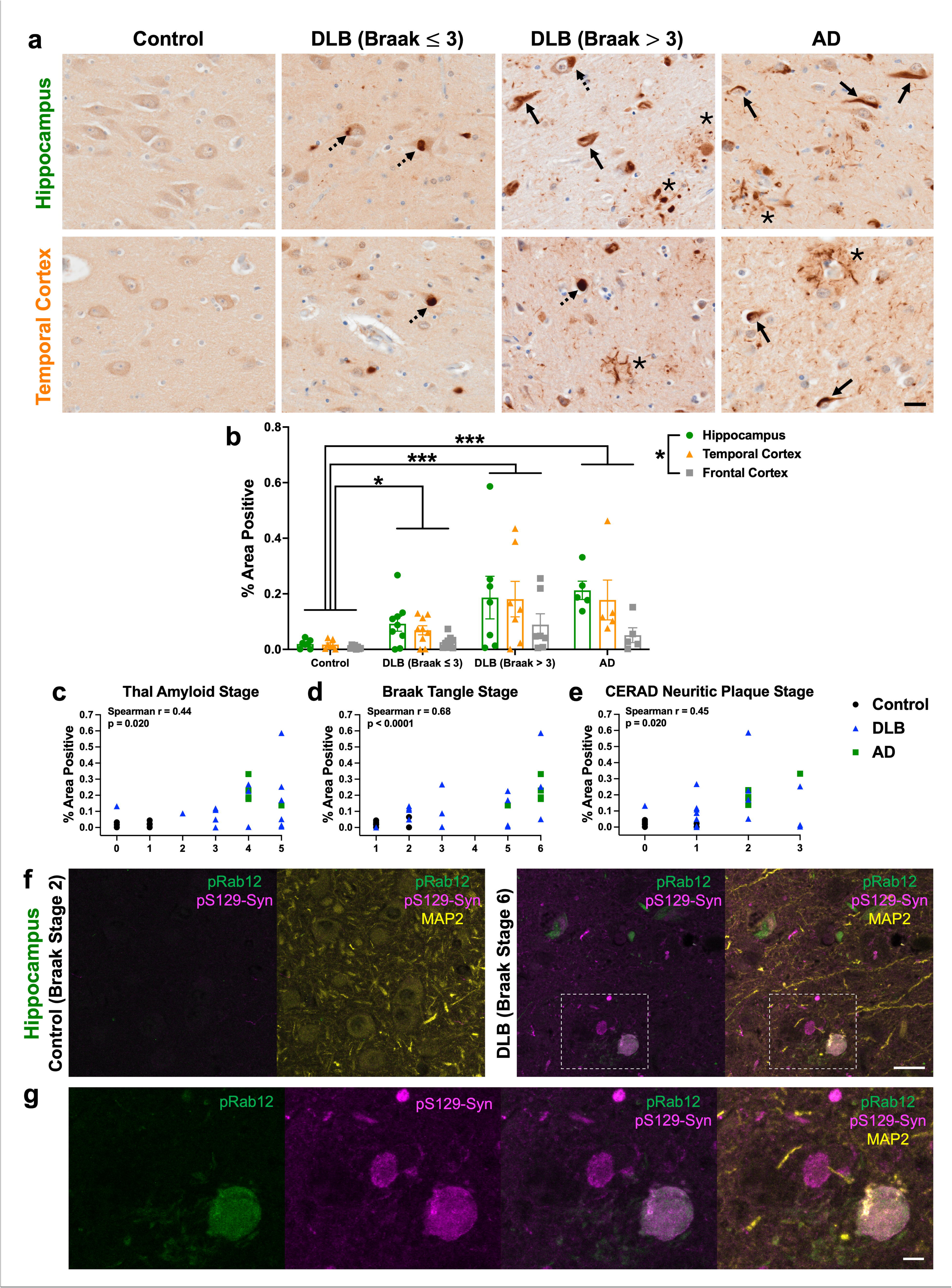
pS106-Rab12 labels tau pathology in DLB and AD and α-synuclein pathology in DLB. (**a**) Representative 60x images of pS106-Rab12 IHC in hippocampus and temporal cortex of a control, DLB (Braak ≤ 3), DLB (Braak > 3), and AD case. Solid arrows denote neurofibrillary tangles, asterisks denote dystrophic neurites, and dashed arrows denote Lewy bodies. (**b**) Quantification of pS106-Rab12-positive area in the hippocampus, temporal cortex and frontal cortex of control (n = 7), DLB (Braak ≤ 3, n = 9), DLB (Braak > 3, n = 7), and AD (n = 5) cases. (**c-e**) Correlations (with Spearman correlation coefficients and associated p-values) of hippocampus pS106-Rab12-positive area with (**c**) Thal amyloid stage, (**d**) Braak tangle stage, and (**e**) CERAD neuritic plaque stage in control, DLB, and AD cases (significant correlations are bolded). (**f**) Representative 63x IF images of pS106-Rab12 (pRab12, green), pS129-Syn (magenta), and MAP2 (yellow) in hippocampus of a control (Braak tangle stage 2) and a DLB subject (Braak tangle stage 6). (**g**) Inset of (**f**) showing co-localization of pRab12 and pS129-Syn in DLB hippocampus. Data are presented as mean ± SEM, and each point represents an individual subject. *p < 0.05, ***p < 0.001 via Kruskal-Wallis H-test with Dunn’s post hoc multiple comparisons test. Scale bars in (**a, f**) = 20µm, scale bar in (**g**) = 5µm

GVBs are associated with early tau pathology, as they are rarely found in neurons with mature tau pathology (i.e., neurofibrillary tangles) [68, 129]. Consistent with this, we infrequently found neurons with pS106-Rab12-labeled GVBs and putative neurofibrillary tangles (Supplementary Fig. 13d), highlighting that pS106-Rab12-labeled GVBs either disappear when neurons reach mature tau tangle stages, or are protective and typically prevent tau from maturing into neurofibrillary tangles.

To demonstrate pS106-Rab12 labels pathological α-synuclein inclusions, we performed IF of pS106-Rab12 and phospho-α-synuclein S129 (pS129-Syn), along with MAP2 as a neuronal marker, in the hippocampus and adjacent temporal cortex of a control subject with Braak tangle stage 2 and a DLB case with Braak tangle stage 6. We found pS106-Rab12 and pS129-Syn signal in the DLB case, but not the control (Fig. 6f). Co-localization of pS106-Rab12 with a subset of Lewy bodies labeled by pS129-Syn was observed (Fig. 6g). Further, we discovered pS106-Rab12 labels a small proportion of pS129-Syn-positive Lewy neurites (Supplementary Fig. 14a), and a subset of pS106-Rab12-labeled GVBs are also positive for pS129-Syn (Supplementary Fig. 14b).

Next, to demonstrate pS106-Rab12 labels tau pathology, we performed IF of pS106-Rab12 and AT8, along with MAP2 as a neuronal marker, in the hippocampus and adjacent temporal cortex of the same subjects. We found pS106-Rab12 and AT8 signal in the DLB case, but not the control (Fig. 7a). pS106-Rab12 GVBs were found in neurons with pre-tangle levels of AT8 labeling, but co-localization of pS106-Rab12 GVBs with AT8 in neurons was not clearly observed (Fig. 7b). Interestingly, we observed co-localization of pS106-Rab12 with a subset of neuropil tau threads labeled by AT8 (Fig. 7b). Further, we discovered pS106-Rab12 labels some, but not all, neurofibrillary tangles (Fig. 7c), and pS106-Rab12 labels, some, but not all, neurites within dystrophic neurites (Fig. 7d). These findings demonstrate 1) pS106-Rab12-labeled GVBs are associated with early, but not late, tau pathology, and 2) pS106-Rab12 localizes to mature tau and α-synuclein pathology.

**Fig. 7.**
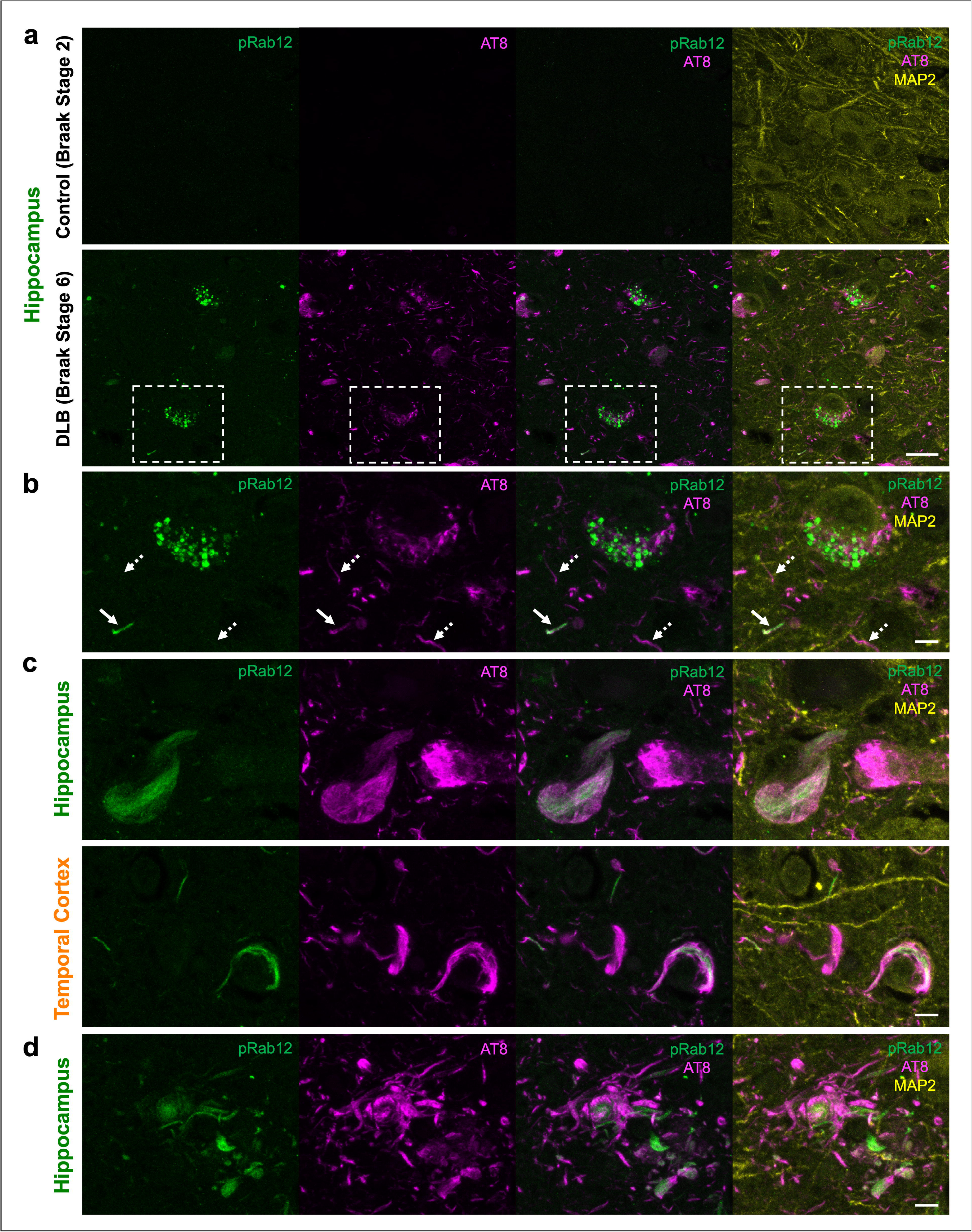
pS106-Rab12 labels a subset of pathological tau inclusions in DLB. (**a**) Representative 63x IF images of pS106-Rab12 (pRab12, green), AT8 (magenta), and MAP2 (yellow) in hippocampus of a control (Braak tangle stage 2) and DLB (Braak tangle stage 6) case. (**b**) Inset of (**a**) showing co-localization of pRab12 GVBs and AT8 in DLB hippocampus. Solid white arrows show a pRab12-positive neuropil tau thread, dashed white arrows show a pRab12-negative neuropil tau thread. (**c**) Representative 63x IF images of pRab12 (green), AT8 (magenta), and MAP2 (yellow) in hippocampus and temporal cortex of a DLB (Braak tangle stage 6) subject showing pRab12-positive and pRab12-negative neurofibrillary tangles. (**d**) Representative 63x images of pRab12 (green), AT8 (magenta), and MAP2 (yellow) IF in hippocampus of a DLB (Braak tangle stage 6) subject showing dystrophic neurites partially labeled by pRab12. Scale bar in (**a**) = 20µm, scale bars in (**b-d**) = 5µm

Similar to AD and DLB, we observed pS106-Rab12 labeling of pathological tau, in the form of neurofibrillary tangles and dystrophic neurites, in the hippocampus and temporal cortex of LRRK2^GS^ PD cases (Braak > 3) and iPD cases (Braak > 3) (Fig. 8a). We also observed pS106-Rab12 labeling of Lewy bodies in LRRK2^GS^ PD and iPD cases (Fig. 8a), although this was not observed in LRRK2^GS^ PD (Braak ≤ 3) cases, likely due to 3 out of 4 of these cases lacking neocortical Lewy bodies (Supplementary Tables 3-4). Consistent with pS106-Rab12 labeling of tau pathology, pS106-Rab12-positive area was significantly elevated in LRRK2^GS^ PD (Braak > 3) compared to both control and LRRK2^GS^ PD (Braak ≤ 3) cases (Fig. 8b). Additionally, pS106-Rab12-positive area was significantly elevated in iPD (Braak > 3), compared to control and LRRK2^GS^ PD (Braak ≤ 3) cases (Fig. 8b). Hippocampus pS106-Rab12-positive area demonstrated a significant positive correlation with Braak neurofibrillary tangle stage, but not Thal amyloid stage, CERAD neuritic plaque stage, age, or PMI (Fig. 8c-e, Supplementary Fig. 15a, b). In contrast, pS106-Rab12 labeling of tau and α-synuclein inclusions was not observed in LRRK2^L1165P^ PD and LRRK2^GS^ Schizophrenia subject brains (Supplementary Fig. 15c).

**Fig. 8.**
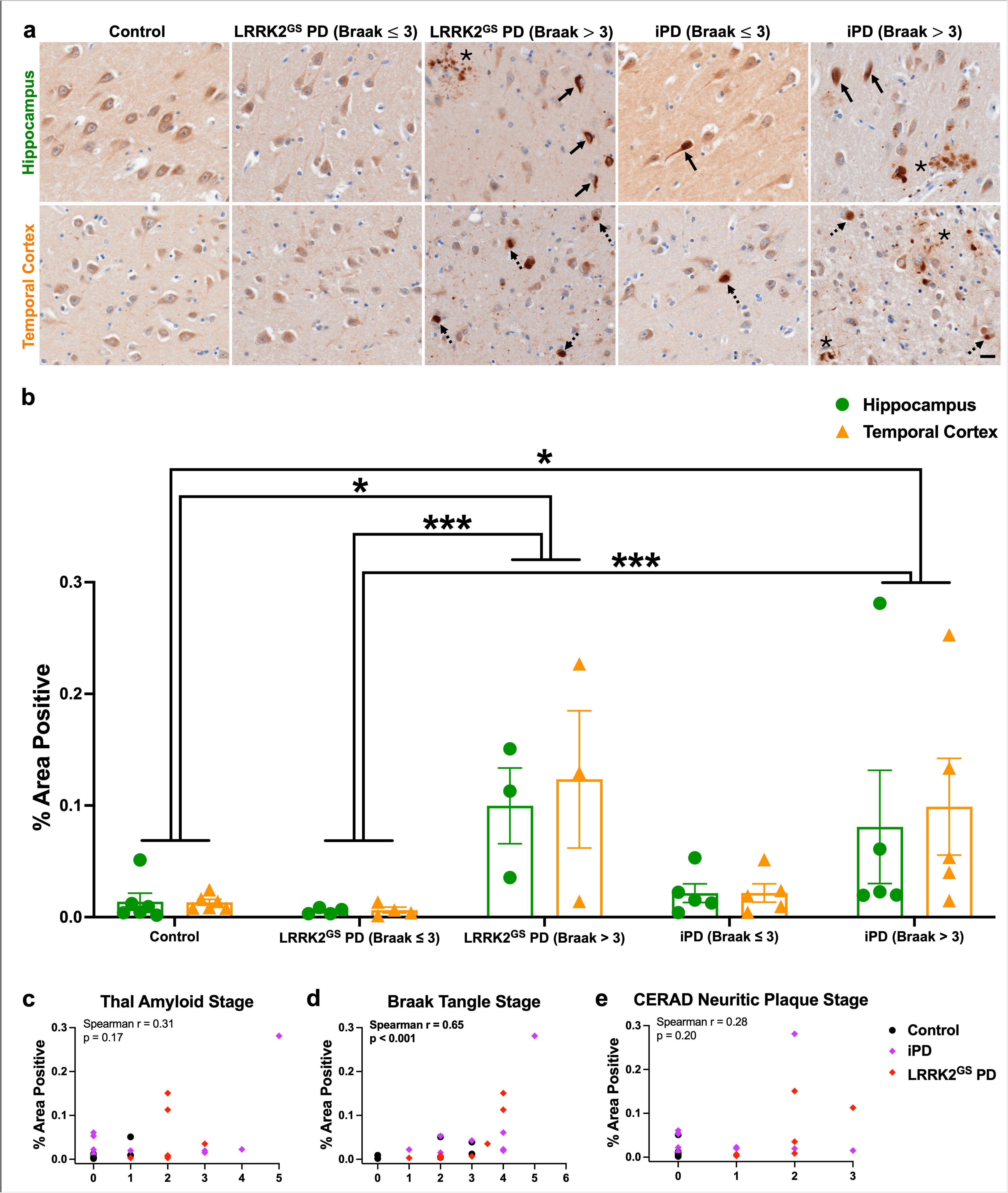
pS106-Rab12 labels tau pathology and Lewy bodies in LRRK2^GS^ PD and iPD. (**a**) Representative 60x images of pS106-Rab12 IHC in hippocampus and temporal cortex of a control, LRRK2^GS^ PD (Braak ≤ 3), LRRK2^GS^ PD (Braak > 3), iPD (Braak ≤ 3), and iPD (Braak > 3) subject. Solid arrows denote neurofibrillary tangles, asterisks denote dystrophic neurites, and dashed arrows denote Lewy bodies. (**b**) Quantification of pS106-Rab12-positive area in the hippocampus of control (n = 6), LRRK2^GS^ PD (Braak ≤ 3, n = 4), LRRK2^GS^ PD (Braak > 3, n = 3), iPD (Braak ≤ 3, n = 5), and iPD (Braak > 3, n = 5) cases. (**c-e**) Correlations (with Spearman correlation coefficients and associated p-values) of hippocampus pS106-Rab12-positive area with (**c**) Thal amyloid stage, (**d**) Braak tangle stage, and (**e**) CERAD neuritic plaque stage in control, LRRK2^GS^ PD and iPD cases (significant correlations are bolded). Data are presented as mean ± SEM, and each point represents an individual subject. *p < 0.05, ***p < 0.001 via Kruskal-Wallis H-test with Dunn’s post hoc multiple comparisons test. Scale bar in (**a**) = 20µm

To determine if LRRK2 localized to these inclusions or GVBs we analyzed IHC of phospho-LRRK2 S935 (pS935-LRRK2), which is not reflective of LRRK2 autophosphorylation activity, in a subset of control, DLB and AD cases (Supplementary Table 1) [26, 53, 65]. Only a cytoplasmic labeling of pS935-LRRK2 was observed, with occasional faint labeling of dystrophic neurites and occasional increased cytoplasmic signal in some DLB and AD cases (Supplementary Fig. 16a). While there were individual differences in pS935-LRRK2 cytoplasmic signal, pS935-LRRK2-positive area was not significantly different across unaffected control, DLB, and AD cases (Supplementary Fig. 16b). These findings suggest that phosphorylated Rab12, but not pS935-LRRK2, is associated with GVBs or mature protein pathology in tauopathies and synucleinopathies.

### pS106-Rab12 labels GVBs and pathological tau inclusions in primary tauopathies

Finally, we sought to determine whether pS106-Rab12 labels pathology in primary tauopathies to demonstrate the direct connection between tau-induced pathological changes and pS106-Rab12 signal. We therefore analyzed pS106-Rab12 IHC in the hippocampus, temporal cortex, frontal cortex, and/or striatum from cases with Pick’s disease, PSP, and CBD (Supplementary Table 1).

We found sparse pS106-Rab12 GVB labeling in hippocampal neurons in PSP and CBD (Fig. 9a), although not in Pick’s disease. This is consistent with previous observations that primary tauopathies display GVBs, although they are sparser than in AD and appear in regions particularly affected by each disease (i.e., they do not follow AD-type GVB staging or progression) [82, 94]. In addition, we observed pS106-Rab12 labeling of putative tau inclusions in all three primary tauopathies. Neuronal labeling of pS106-Rab12 in the Dentate Gyrus of the hippocampus in Pick’s disease, consistent with Pick bodies, as well as neuronal labeling of potential tau inclusions in other hippocampal regions of Pick’s disease was observed (Fig. 9b). We also found occasional pS106-Rab12 labeling of neurofibrillary tangles in the temporal cortex of PSP cases, and light labeling of ballooned neurons in the frontal cortex and striatum of CBD cases (Fig. 9b). Finally, pS106-Rab12-positive glial inclusions were observed in the temporal cortex of Pick’s disease, and the hippocampus, temporal cortex and frontal cortex of PSP and CBD (Fig. 9c, Supplementary Fig. 17). This labeling may reflect high glial tau pathology in primary tauopathies [14]. The pattern of labeling does not clearly reflect PSP tufted astrocytes or CBD astrocytic plaques, rather it shows closer resemblance to oligodendroglial tau inclusions like coiled bodies. In all, these findings demonstrate pS106-Rab12 labels pathological inclusions in neurodegenerative diseases that are primarily driven by pathological tau accumulation.

**Fig. 9.**
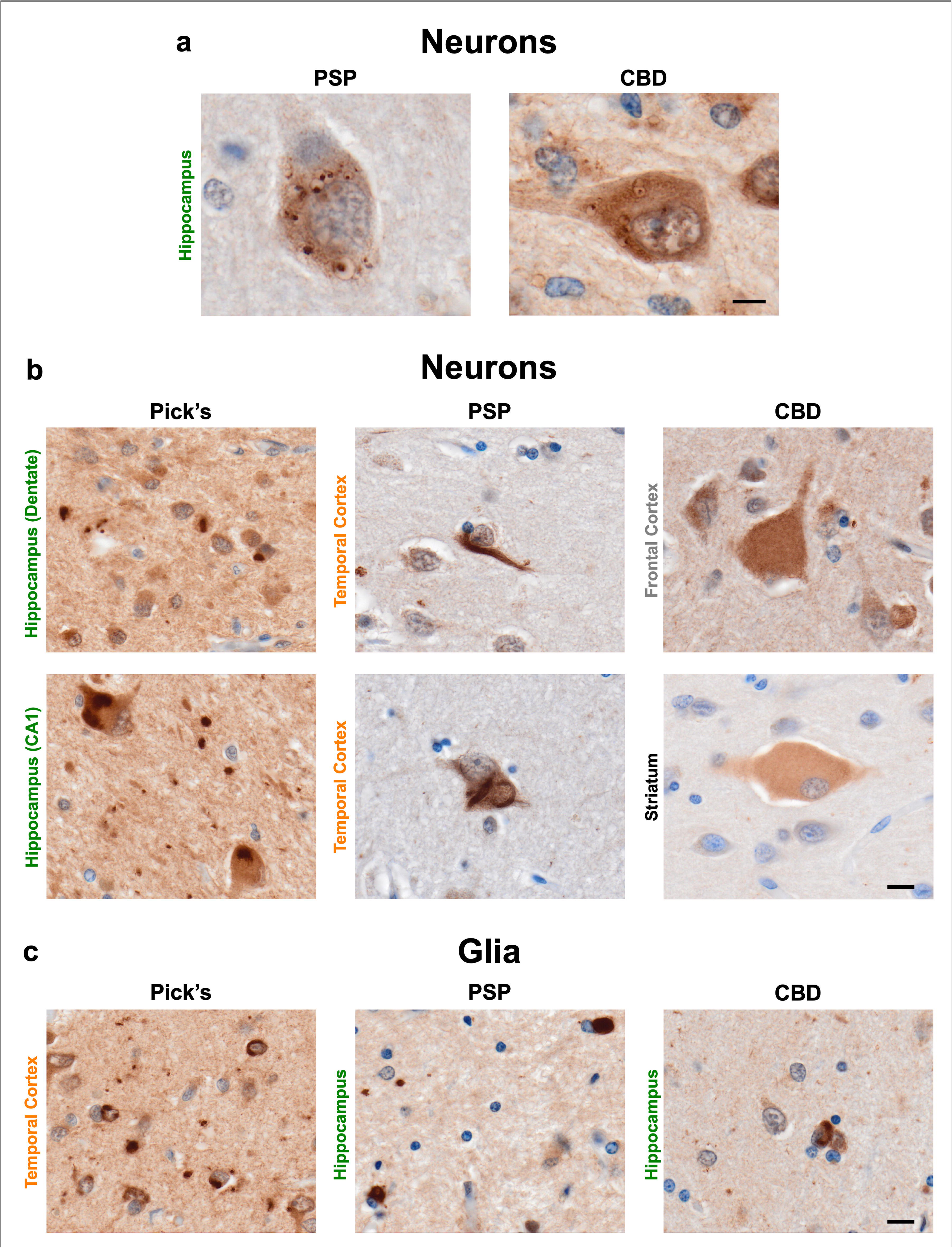
pS106-Rab12 labeling in primary tauopathies. (**a**) Representative 60x images of pS106-Rab12-labeled GVBs in hippocampus neurons from PSP and CBD cases. (**b**) Representative 60x images of pS106-Rab12 labeling of putative tau pathology in hippocampus, temporal cortex, frontal cortex, and/or striatum neurons from Pick’s disease (n = 3), PSP (n = 3) and CBD cases (n = 3). (**c**) Representative 60x images of pS106-Rab12 labeling in temporal cortex or hippocampus glial cells from Pick’s disease, PSP and CBD cases. Scale bar in (**a**) = 5µm, scale bars in (**b, c**) = 10µm

## Discussion

There is emerging evidence that elevated LRRK2 kinase activity is observed not just in LRRK2 PD, but in non-genetic or idiopathic forms of PD. While there are many proposed mechanisms of enhancing LRRK2-mediated Rab substrate phosphorylation, a major activator is endolysosomal stress. Herein we describe that LRRK2-mediated phosphorylated Rab12 labels lysosomal structures, GVBs, in *LRRK2* mutation carriers and idiopathic PD – and across clinically distinct neurodegenerative diseases (AD, DLB and primary tauopathies) not previously reported to be characterized by LRRK2 pathway dysfunction. In addition to labeling of these lysosomal GVBs, phosphorylated Rab12 was increased in the hippocampus and temporal cortex and labeled a subset of mature pathological tau and α-synuclein inclusions in human tauopathies and synucleinopathies. pS106-Rab12 labeling of GVBs, which are closely associated with early pathological tau accumulation, was recapitulated in the PS19 mouse tauopathy model. Overall, we have identified a potential link between lysosomes, LRRK2 kinase activity and neuronal vulnerability to pathological protein accumulation across neurodegenerative diseases.

In addition to being causative for LRRK2 PD, *LRRK2* variants increase risk for development of iPD and the primary tauopathy PSP [6, 39, 63, 64, 79, 83, 99]. Recent genetic evidence suggests *LRRK2* mutations may also play a role in development of DLB, particularly in people with North African ancestry [31, 73]. Further, single nucleotide polymorphisms in the gene encoding Rab10 protein, a LRRK2 substrate, are associated with risk and resilience to AD development [49, 67]. Our findings of increased Rab12 phosphorylation and pS106-Rab12 labeling of GVBs and pathology across synucleinopathies and tauopathies suggest that LRRK2 plays a role in other neurodegenerative diseases in addition to Parkinsonism. Given that our findings indicate increased Rab12 phosphorylation in the brain in multiple neurodegenerative diseases, next steps may include investigating whether peripheral pS106-Rab12, which was found to be elevated in LRRK2^GS^ PD patients and LRRK2^GS^ non-manifesting carriers compared to controls, is similarly increased and extends beyond PD using a variety of these approaches [15, 97].

Since the identification of increased kinase activity associated with *LRRK2* pathogenic mutations [93], it has been difficult to reliably resolve LRRK2 activity-dependent markers in animal models and post-mortem tissue banks. LRRK2 autophosphorylation at S1292 directly reflects LRRK2 kinase activity, but low endogenous levels in tissue and other technical issues present barriers [20, 28, 46, 74]. LRRK2 kinase inhibition via type I kinase inhibitors leads to LRRK2 dephosphorylation at S935, and is therefore considered an indirect marker of LRRK2 kinase activity [26, 53, 65]. However, measurements of pS935-LRRK2 in human brain, peripheral blood neutrophils, lymphoblastoid cell lines and peripheral blood mononuclear cells in several studies across multiple laboratories demonstrate that pS935-LRRK2 levels, relative to total LRRK2, are either unchanged or even decreased in LRRK2 PD and iPD cases compared to controls, and sometimes decreased in LRRK2 PD compared to iPD cases [25–27, 62, 65, 88, 92]. One study found no change in pS935-LRRK2 protein relative to β-actin in the temporal cortex of DLB cases compared to controls [85]. Consistent with the above findings showing no change or even decreased pS935-LRRK2 in PD and DLB, we found no significant difference in pS935-LRRK2 in the brains from DLB or AD cases compared to control brains, and as such does not appear to be elevated in neurodegenerative disease brain tissue. Though, our observation of diffuse cytoplasmic labeling of pS935-LRRK2 in human neurons is consistent with previous work [25].

We found that pS106-Rab12 labels pathological tau inclusions. While interactome studies of tau pathology do not show enrichment of Rab12 in tau inclusions or neurons with tau inclusions, studies have identified LRRK2 and other LRRK2-associated proteins as tau interactors [21, 58, 59, 98]. Probe-dependent proximity profiling of tau pathology found an association of vacuolar protein sorting-associated protein 35 (VPS35) with tau pathology across tauopathies [59]. The D620N mutation in *VPS35* causes autosomal dominant PD and increases LRRK2 kinase activity [58, 98]. Two interactome studies have also identified Rab10 as an interactor with tau inclusions across tauopathies [21, 59]. While these findings implicate LRRK2 pathway proteins as pathological tau interactors, Rab12 may not have been identified because only phosphorylated Rab12 is specifically enriched at tau inclusions. Future phospho-proteomic interactome work should investigate this possibility.

We observed elevated Rab12 phosphorylation in hippocampus and entorhinal cortex of DLB cases, as well as increased pS106-Rab12 labeling across neurodegenerative diseases with tau and/or α-synuclein pathology. In mice, Rab12 phosphorylation is reliably detected in the brain and reflects levels of LRRK2 kinase activity [44, 47, 58]. Further, a recent study shows elevated pS106-Rab12 labeling in cholinergic neurons in aged LRRK2^GS^ knock-in mice compared to wild-type [11]. This elevation in pS106-Rab12 labeling is consistent with our finding of age-dependent accumulation of pS106-Rab12-positive GVBs in PS19 mice as well as other work showing age-dependent accumulation of GVBs in mouse tauopathy and synucleinopathy models [22, 36, 42, 95]. Elevated pS106-Rab12 labeling in cholinergic neurons may reflect vulnerability of cholinergic regions, like the nucleus basalis of Meynert and pedunculopontine tegmentum, to GVB accumulation [82].

GVBs are lysosomal structures, and tau and α-synuclein aggregation both induce endolysosomal stress and can be degraded by lysosomes [12, 34, 41, 42, 48, 50, 54, 66, 68, 69, 77, 95]. While the mechanism underlying GVB formation is unknown, the fact that GVBs are associated with early, but not late, tau pathology suggests that GVBs may be either damaging (contributing to progression or maturation of aggregates) or protective (preventing mature aggregates from forming, while neurons without GVBs develop mature tangles/Lewy bodies) [94]. One limitation to this study is that it is unknown whether pS106-Rab12 localization to GVBs contributes to the hypothesized damaging effect of GVBs (by inducing their formation) or reduces the potential protective effect of GVBs (by impairing their degradative capacity). Future work therefore may investigate both the role of pS106-Rab12 in mediating GVB formation as well as the protective or damaging effect of GVBs within neurons. Additionally, the mechanism(s) by which tau and/or α-synuclein pathology mediates pS106-Rab12 accumulation, or how LRRK2 activity mediates tau and/or α-synuclein aggregation, are unclear. Previous findings are conflicting with regards to the effect of elevated LRRK2 kinase activity on tau and α-synuclein aggregation, reporting either an exacerbation of pathological protein accumulation by pathogenic *LRRK2* mutation or no effect of pathogenic *LRRK2* [2, 18, 23, 24, 52, 56, 60, 71, 89]. It is also unknown why pS106-Rab12 labels a subset of tau and α-synuclein pathological inclusions. pS106-Rab12 may label a subset because some neurons with inclusions develop GVBs, but not others. Understanding differences between pS106-Rab12-positive and pS106-Rab12-negative protein inclusions will be important for targeting LRRK2 in PD.

In summary, we found increased Rab12 phosphorylation and localization of pS106-Rab12 to tau and α-synuclein pathology in tauopathies and synucleinopathies, including LRRK2 PD, iPD, DLB, AD and primary tauopathies. We also discovered pS106-Rab12 labels GVBs which were initiated by pathological tau accumulation. These findings associate LRRK2 kinase activity and the LRRK2 substrate Rab12 to neurodegenerative diseases with tau and/or α-synuclein pathology. Future work will investigate mechanisms by which the LRRK2 pathway associates with and mediates protein aggregation and GVBs, which may uncover interventions targeting protein aggregation related to LRRK2 activity, both in LRRK2 PD and across tauopathies and synucleinopathies.

## Supporting information

Buck et al Supplementary Figures

Buck et al Supplementary Tables

## List of abbreviations

AD: Alzheimer’s disease
Ck1δ: casein kinase 1δ
CBD: corticobasal degeneration
DLB: Dementia with Lewy bodies
GVB: granulovacuolar degeneration body
iPD: idiopathic Parkinson’s disease
KO: knockout
LAMP1: lysosomal-associated membrane protein 1
LRRK2: leucine-rich repeat kinase 2
LRRK2^GS^: LRRK2 G2019S
PD: Parkinson’s disease
PPase: phosphatase
pS129-Syn: phospho-α-synuclein S129
pS935-LRRK2: phospho-LRRK2 S935
pS106-Rab12/pRab12: phospho-Rab12 S106
pS65-Ub: phospho-Ubiquitin S65
PMI: post-mortem interval
PSP: progressive supranuclear palsy
WT: wild-type

## Acknowledgements

We thank the patients and their families for participating in and contributing to these research studies and their generous donation of brain samples for research. We also thank the Bryan Brain Bank and Biorepository of the Duke-UNC Alzheimer’s Disease Research Center (ADRC), University of Washington Biorepository and Integrated Neuropathology Laboratory, and the University of Pennsylvania Center for Neurodegenerative Disease Research Brain Bank for processing and sharing human autopsy brain tissue. We thank the Duke Light Microscopy Core Facility (LMCF) for microscope access and assistance.

## Authors’ contributions

L.H.S. conceptualized and supervised the research. S.A.B. performed the majority of experiments and contributed to research conception, design and interpretation. T.J.C. and A.B.W. also contributed to interpretation. S.H.J.W. provided human brain tissue, neuropathology diagnosis and analysis, as well as helped interpret human IHC results. J.E. performed human DAB IHC experiments. E.B.M. performed and analyzed mouse western blot experiments. S.A.B. performed human IF experiments and imaging of all human and mouse IHC/IF experiments. S.A.B. analyzed human IHC and IF images. S.A.B. and S.Y. performed mouse IF, and S.A.B., S.Y. and H.W.P. analyzed mouse IF images. T.M. performed and analyzed human western blot experiments. S.A.B. and L.H.S. wrote the manuscript with input from all authors. All authors contributed to reviewing the final version of the manuscript.

## Funding

This work was supported by National Institutes of Health (NIH) R01NS119528 to L.H.S., NIH R21AG084216 to L.H.S. and T.J.C., NIH R01NS064934 to A.B.W., Parkinson’s Foundation PF-PRF-1244721 to S.A.B., NIH P30AG072958 to the Duke-UNC Alzheimer’s Disease Research Center (ADRC), NIH P30AG066509 and NIH U19AG066567 to the University of Washington, and NIH P30AG072979, NIH P01AG066597, and NIH P01AG084497 to the University of Pennsylvania Center for Neurodegenerative Disease Research Brain Bank.

### Data availability

The data used for this study is available from the corresponding author upon request.

## Declarations

### Ethics approval and informed consent

All donors had provided written informed consent for the use of autopsy material and of clinical and genetic information for research purposes. All animal protocols were approved by Duke University’s Institutional Animal Care and Use Committee.

