## Supplementary material for "Accumulation of LRRK2-associated phospho-Rab12 degenerative lysosomes in tauopathies": Buck et al Supplementary Figures

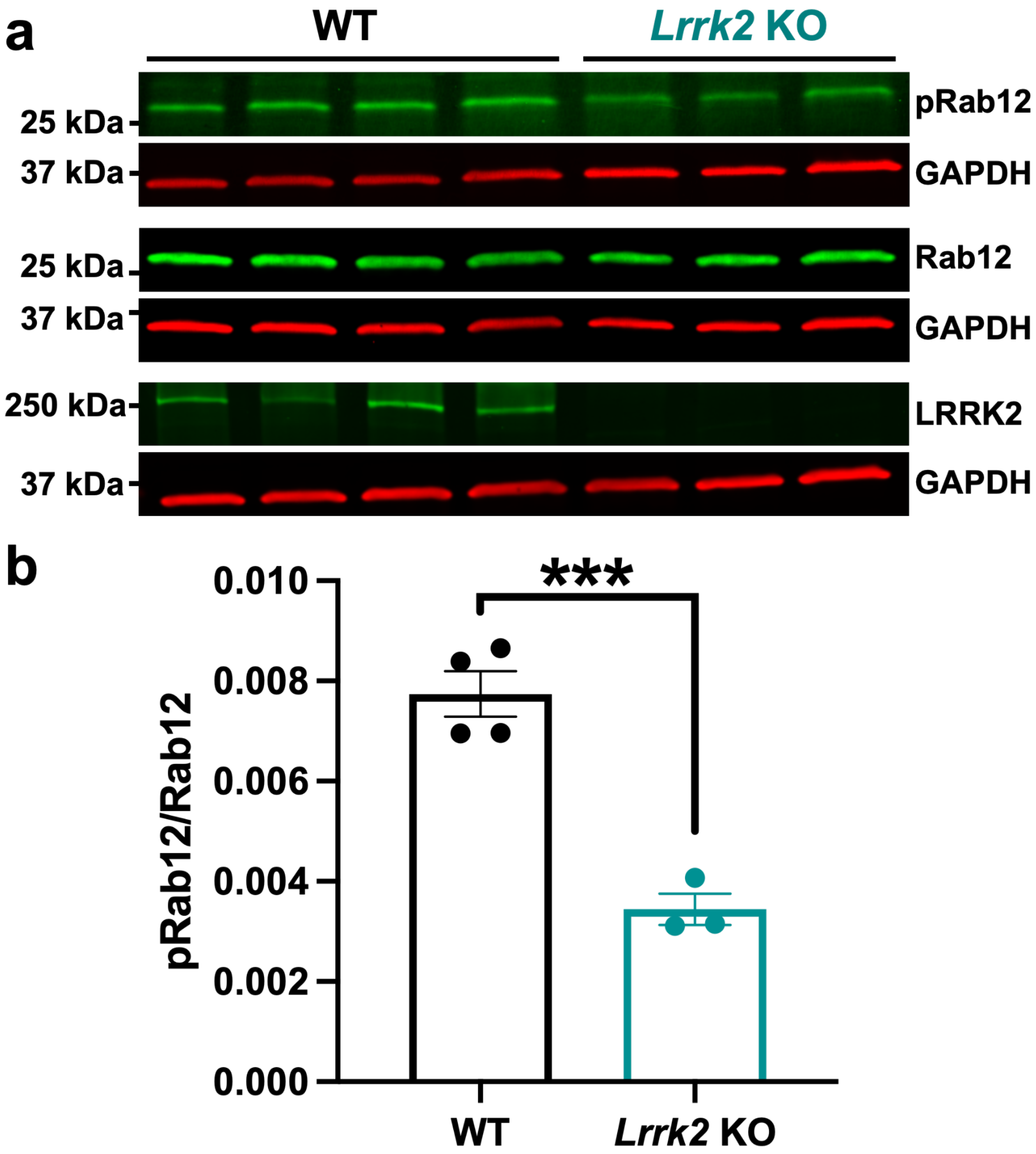


**Supplementary Fig. 1. LRRK2-dependent Rab12 phosphorylation in mice.** (**a**) Representative western blot of cortex derived from WT (n = 4) and *Lrrk2* KO (n = 3) mice assessed for pS106-Rab12 (pRab12), Rab12, LRRK2, and GAPDH as a loading control. (**b**) Quantification of western blots show that pRab12 levels in *Lrrk2* KO mice are significantly decreased 55% compared to WT. Data is presented as pRab12/Rab12 (for which both pRab12 and Rab12 were first normalized to GAPDH). ***$p < 0.001$, determined by unpaired Student's *t* test. Data are presented as mean ± SEM, and each point represents an individual animal.

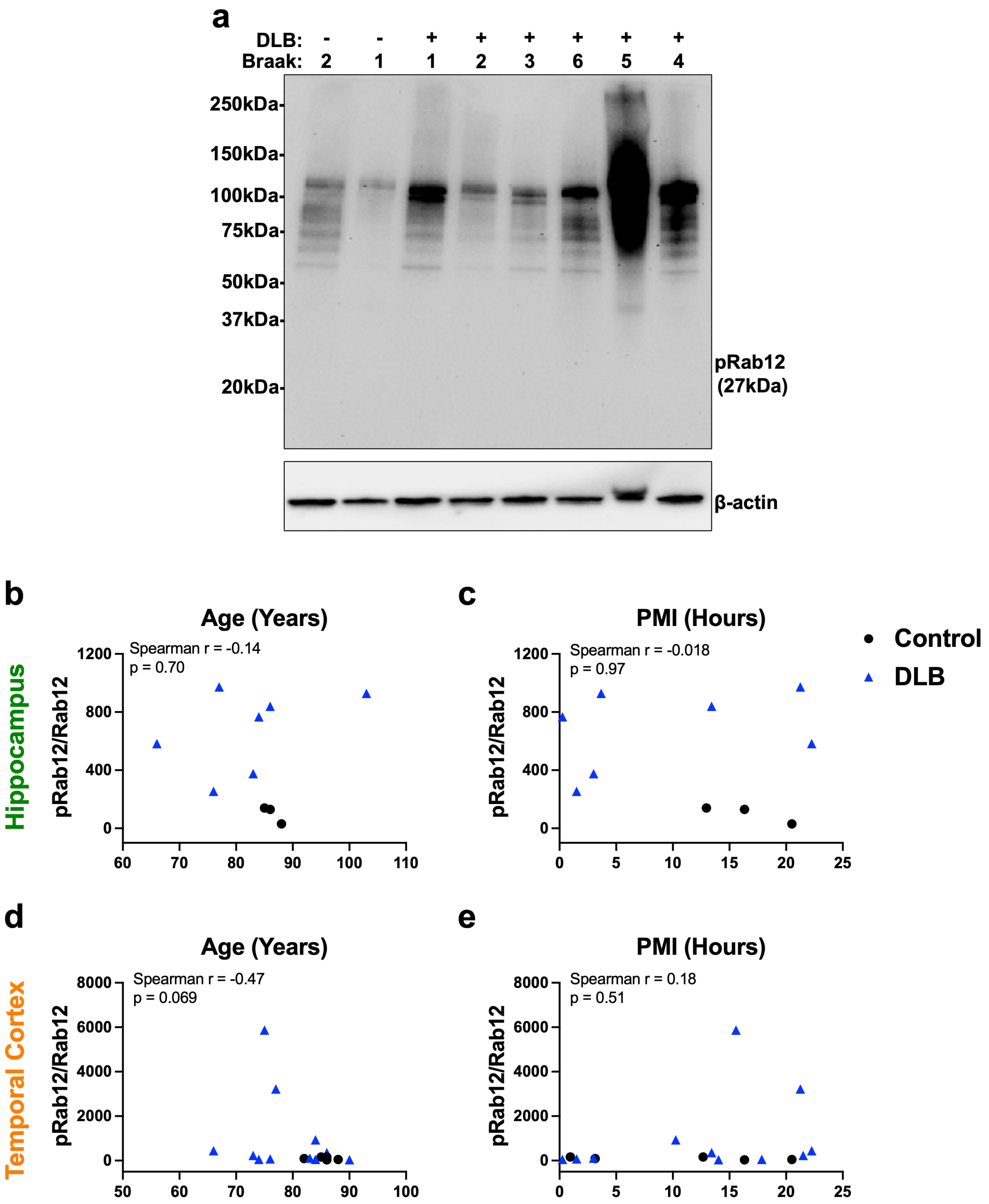


**Supplementary Fig. 2. Western blot analysis for insoluble pS106-Rab12 and correlations of soluble pS106-Rab12 with age and PMI in brains derived from control and DLB cases.**

(**a**) Representative western blot of temporal cortex derived from control (n = 2) and DLB (n = 6) cases for insoluble pS106-Rab12 (pRab12) and β-actin as a loading control. (**b**-**e**) Correlations (with Spearman correlation coefficients and associated p-values) of (**b, c**) hippocampus and (**d, e**) temporal cortex of soluble pRab12/Rab12 with (**b, d**) age and (**c, e**) PMI in control (hippocampus n = 3, temporal cortex n = 5) and DLB (hippocampus n = 7, temporal cortex = 11) cases. Each point represents an individual subject.

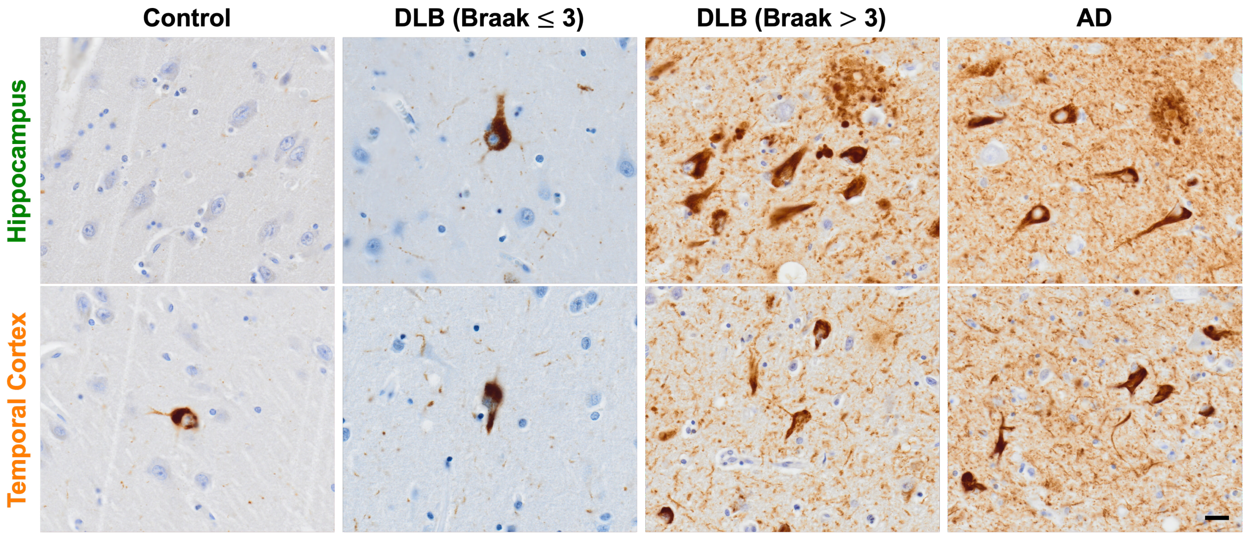


**Supplementary Fig. 3. Representative 60x images of AT8-labeled tau in hippocampus and temporal cortex of a control, DLB (Braak ≤ 3), DLB (Braak > 3), and AD subject.** Scale bar = 20μm.

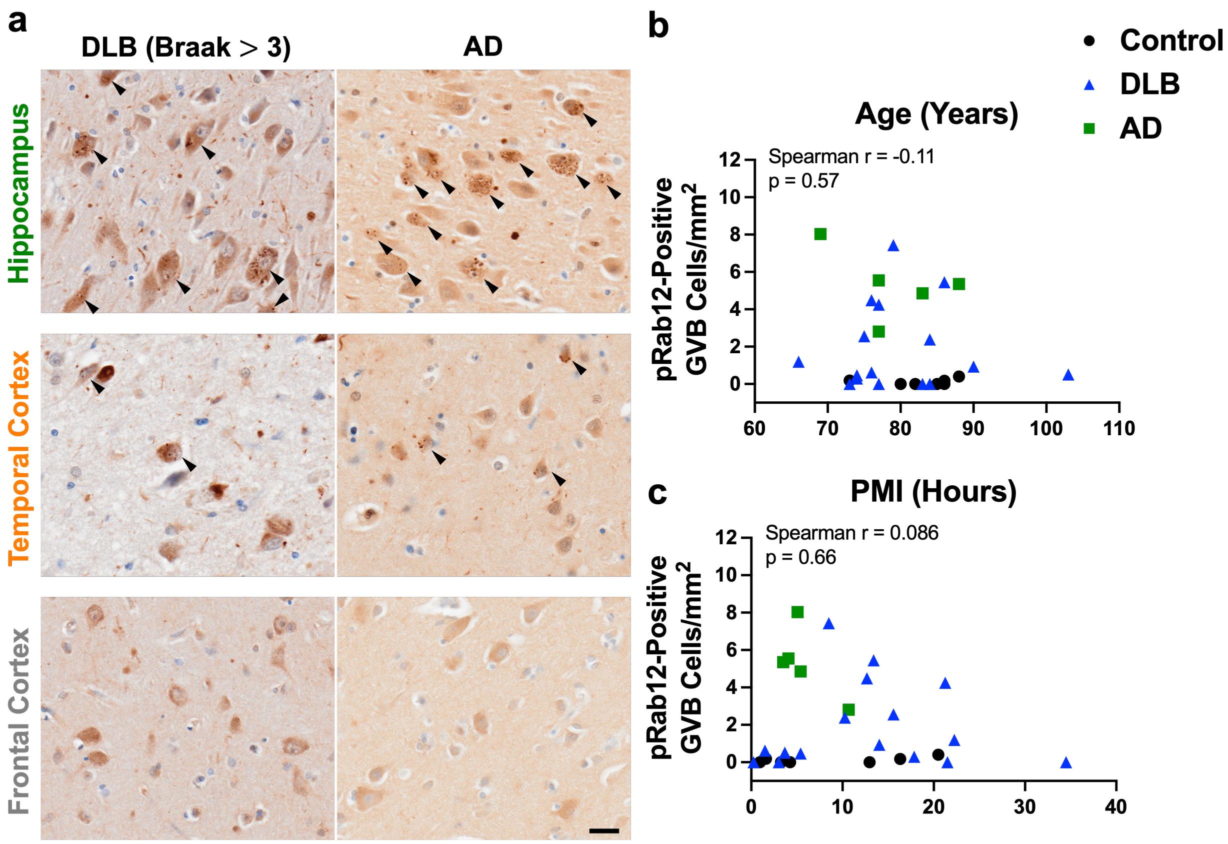


**Supplementary Fig. 4. Brain regional differences in pS106-Rab12 GVB labeling and correlations of Rab12 phosphorylation-linked GVB cell density with age and PMI in control, DLB and AD subjects.** (**a**) Representative 60x images of pS106-Rab12 labeling of GVBs in hippocampus, temporal cortex, and frontal cortex of DLB (Braak > 3) and AD subjects. Arrows denote pS106-Rab12 GVB-positive cells. (**b, c**) Correlations (with Spearman correlation coefficients and associated p-values) of hippocampus pS106-Rab12 GVB cell density with (**b**) age and (**c**) PMI in control (n = 7), DLB (n = 16) and AD (n = 5) cases. Each point represents an individual subject. Scale bar = 20μm.

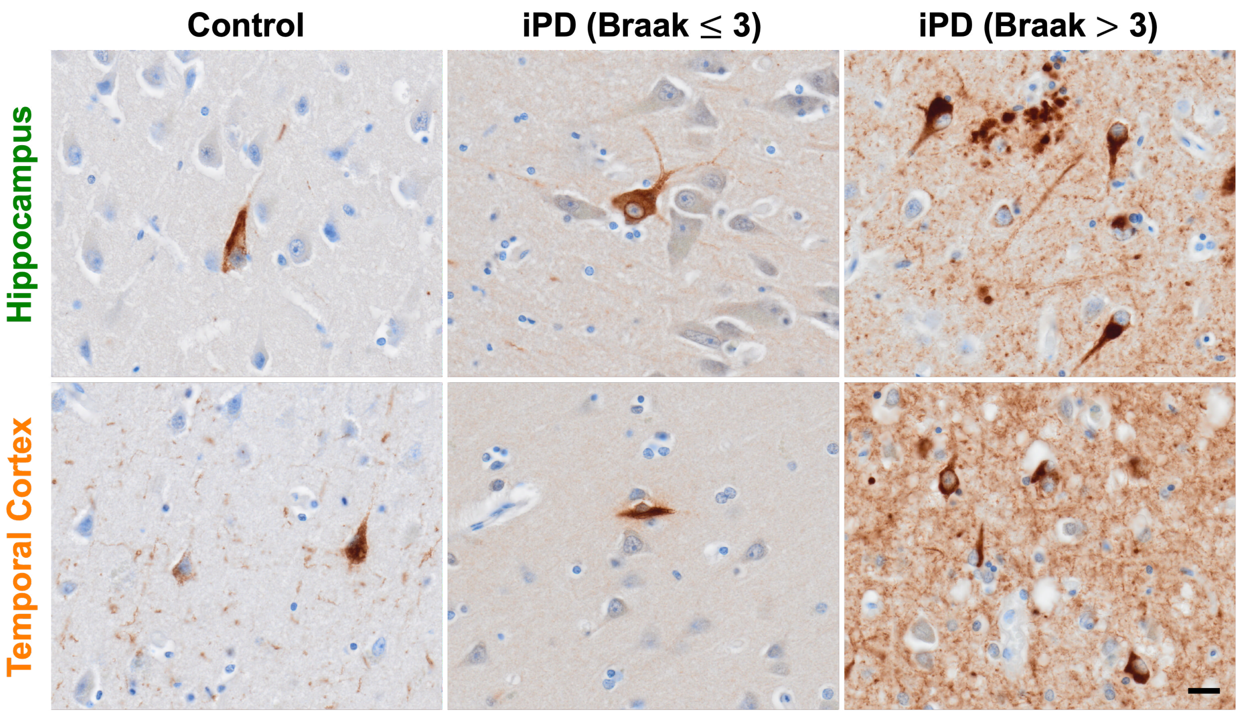


**Supplementary Fig. 5. Representative 60x images of AT8-labeled tau in hippocampus and temporal cortex of a control, iPD (Braak ≤ 3), and iPD (Braak > 3) case.** Scale bar = 20μm.

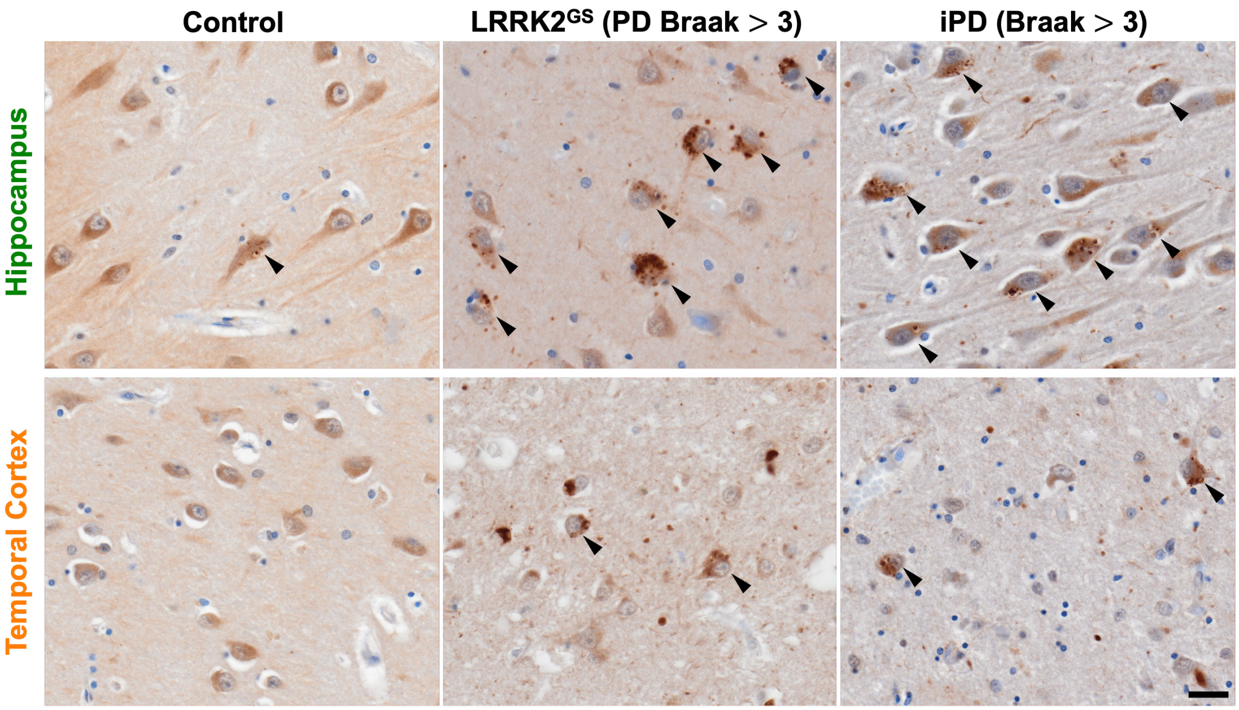


**Supplementary Fig. 6. Regional difference in pS106-Rab12 GVB labeling in control, LRRK2$^{GS}$ PD and iPD cases.** Representative 60x images of pS106-Rab12 labeling of GVBs in hippocampus and temporal cortex of a control, LRRK2$^{GS}$ PD (Braak > 3) and iPD (Braak > 3) case. Arrows denote pS106-Rab12 GVB-positive cells. Scale bar = 20µm.

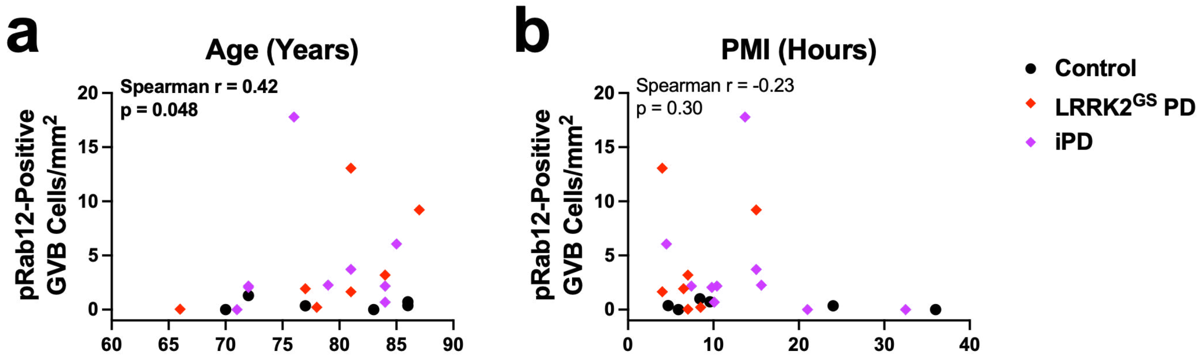


**Supplementary Fig. 7. Correlation of pS106-Rab12 GVB cell density with age and PMI in control, LRRK2$^{GS}$ PD and iPD cases.** (**a**, **b**) Correlations (with Spearman correlation coefficients and associated p-values) of hippocampus pS106-Rab12 GVB cell density with (**a**) age and (**b**) PMI in control (n = 6), LRRK2$^{GS}$ PD (n = 7) and iPD (n = 10) cases (significant correlations are bolded). Each point represents an individual subject.

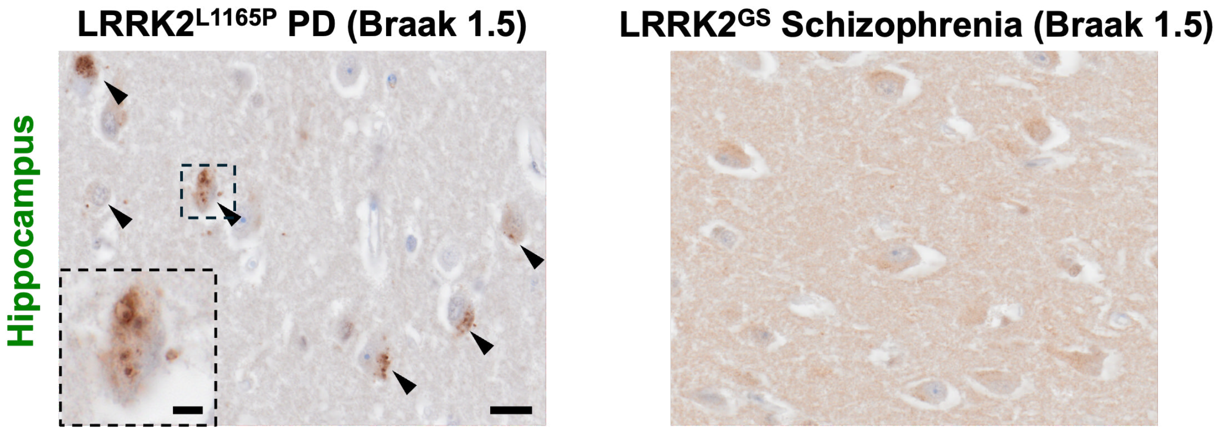


**Supplementary Fig. 8. pS106-Rab12 GVB labeling in LRRK2$^{L1165P}$ PD, but not LRRK2$^{GS}$ Schizophrenia case.** Representative 60x images of hippocampus pS106-Rab12 labeling of GVBs in a LRRK2$^{L1165P}$ PD and a LRRK2$^{GS}$ Schizophrenia case. pS106-Rab12-labeled GVBs were observed in the LRRK2$^{L1165P}$ PD (inset), but not the LRRK2$^{GS}$ Schizophrenia case. Scale bar = 20μm, inset scale bar in = 5μm.

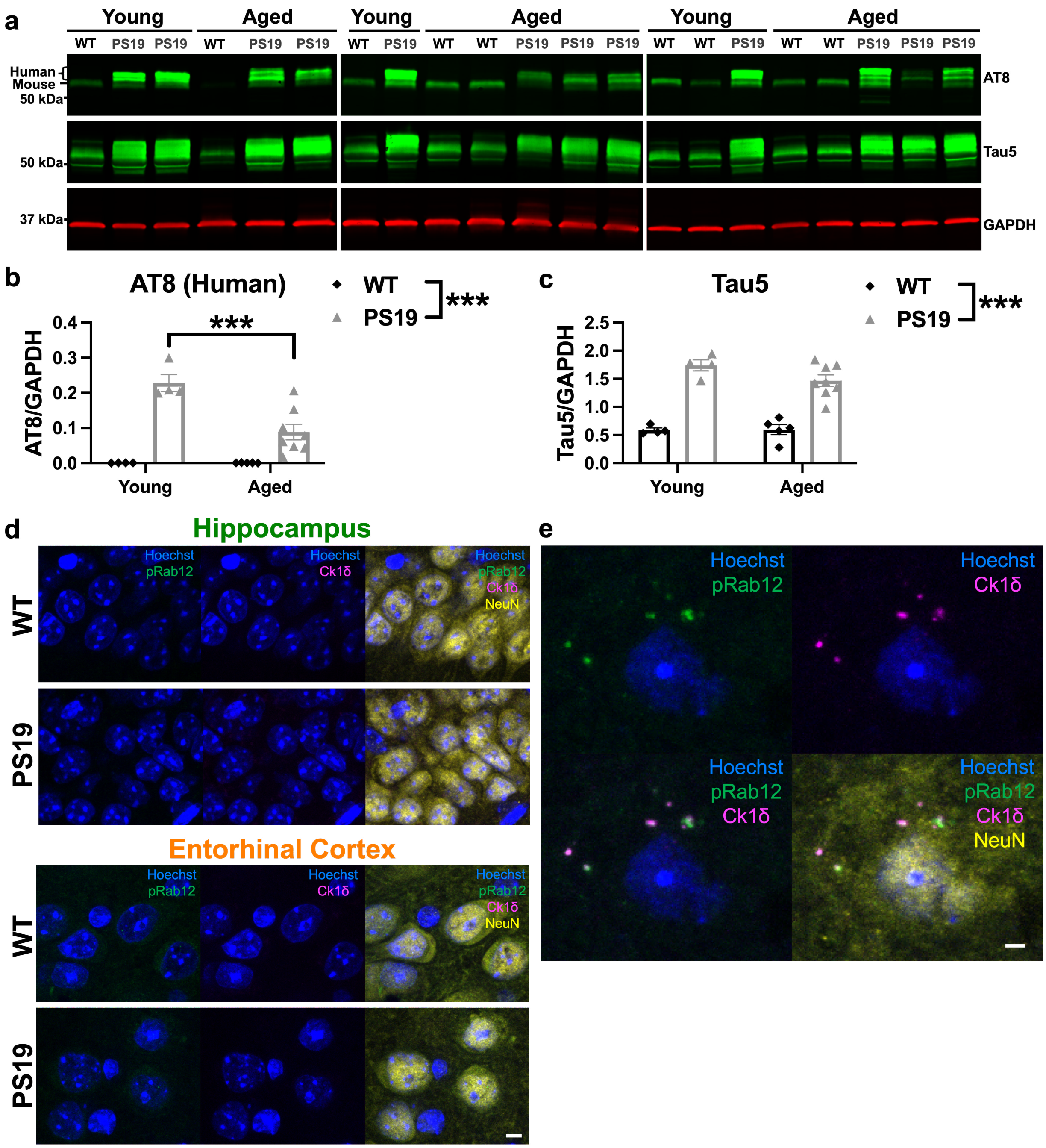


**Supplementary Fig. 9. Increased phosphorylated and total tau in PS19 mouse entorhinal cortex, and lack of pS106-Rab12-labeled GVBs in young PS19 mouse entorhinal cortex.** (**a**) Representative western blots of entorhinal cortex derived from young WT (n = 4), young PS19 (n = 4), aged WT (n = 5) and aged PS19 (n = 8) mice assessed for AT8, Tau5 (total tau), and GAPDH as a loading control. (**b, c**) Quantification of (**b**) AT8/GAPDH (human bands) and (**c**) Tau5/GAPDH show increased AT8 and Tau5 in PS19 entorhinal cortices compared to WT. (**d**) Representative 63x images of Hoechst (blue), pS106-Rab12 (pRab12, green), Ck1δ (magenta), and NeuN (yellow) in hippocampus and entorhinal cortex of young WT and PS19 mice. (**e**) Zoomed-in 63x representative images of young PS19 entorhinal cortex showing overlap of pRab12 and Ck1δ puncta in a GVB-positive cell observed at this age. Data are presented as mean ± SEM, and each point represents an individual animal. ***$p < 0.001$ via two-way ANOVA with Bonferroni's post hoc multiple comparisons test. Scale bar in (**d**) = 5μm, scale bar in (**e**) = 2μm.

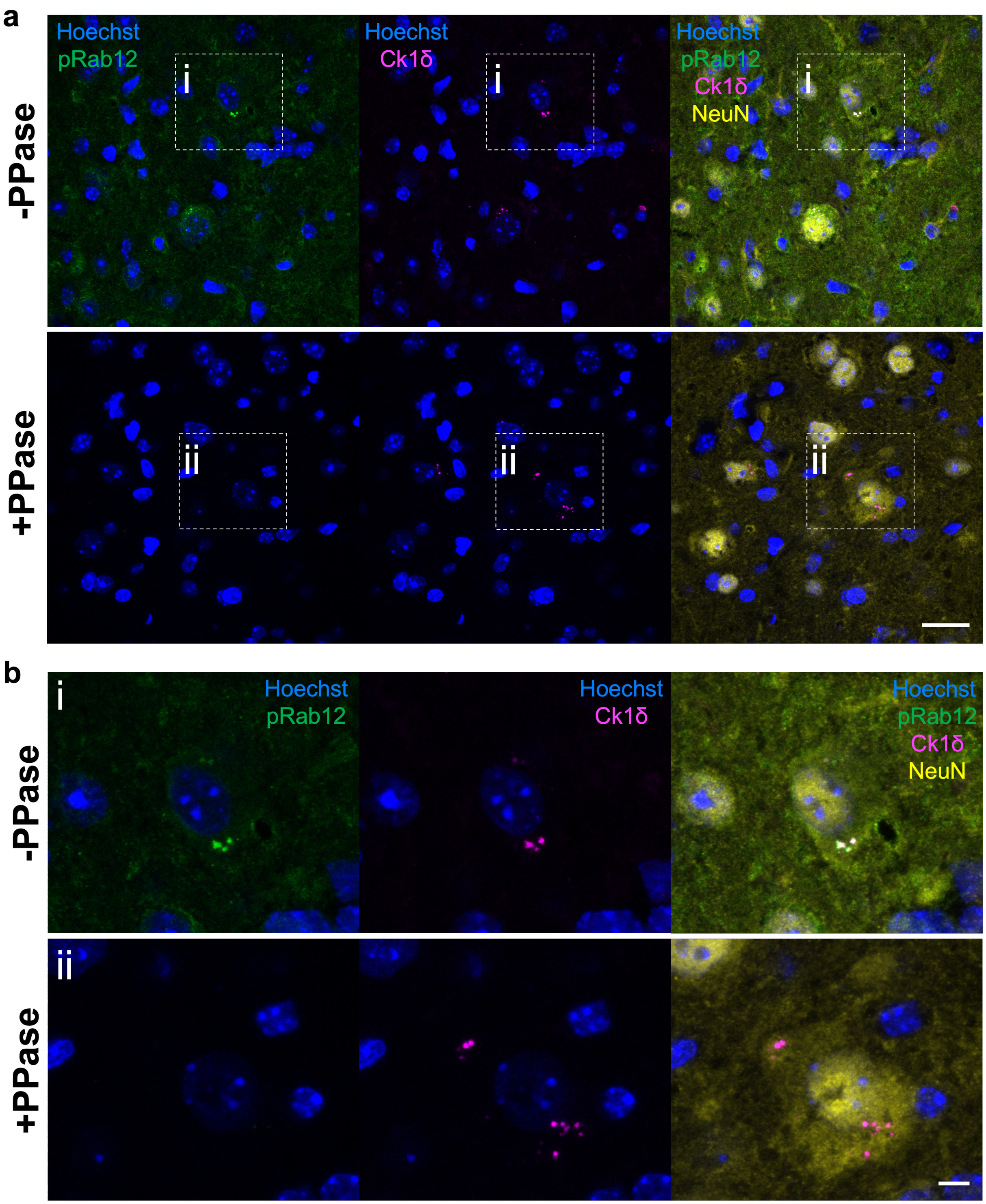


**Supplementary Fig. 10. Phosphatase (PPase) pre-treatment abolishes pS106-Rab12 GVB signal in aged PS19 mouse entorhinal cortex.** (**a**) Representative 63x IF images and (**b**) insets of Hoechst (blue), pS106-Rab12 (pRab12, green), Ck1δ (magenta), and NeuN (yellow) in aged

PS19 mouse entorhinal cortex with (+) or without (-) PPase pre-treatment. Scale bar in (**a**) = 20μm, scale bar in (**b**) = 5μm.

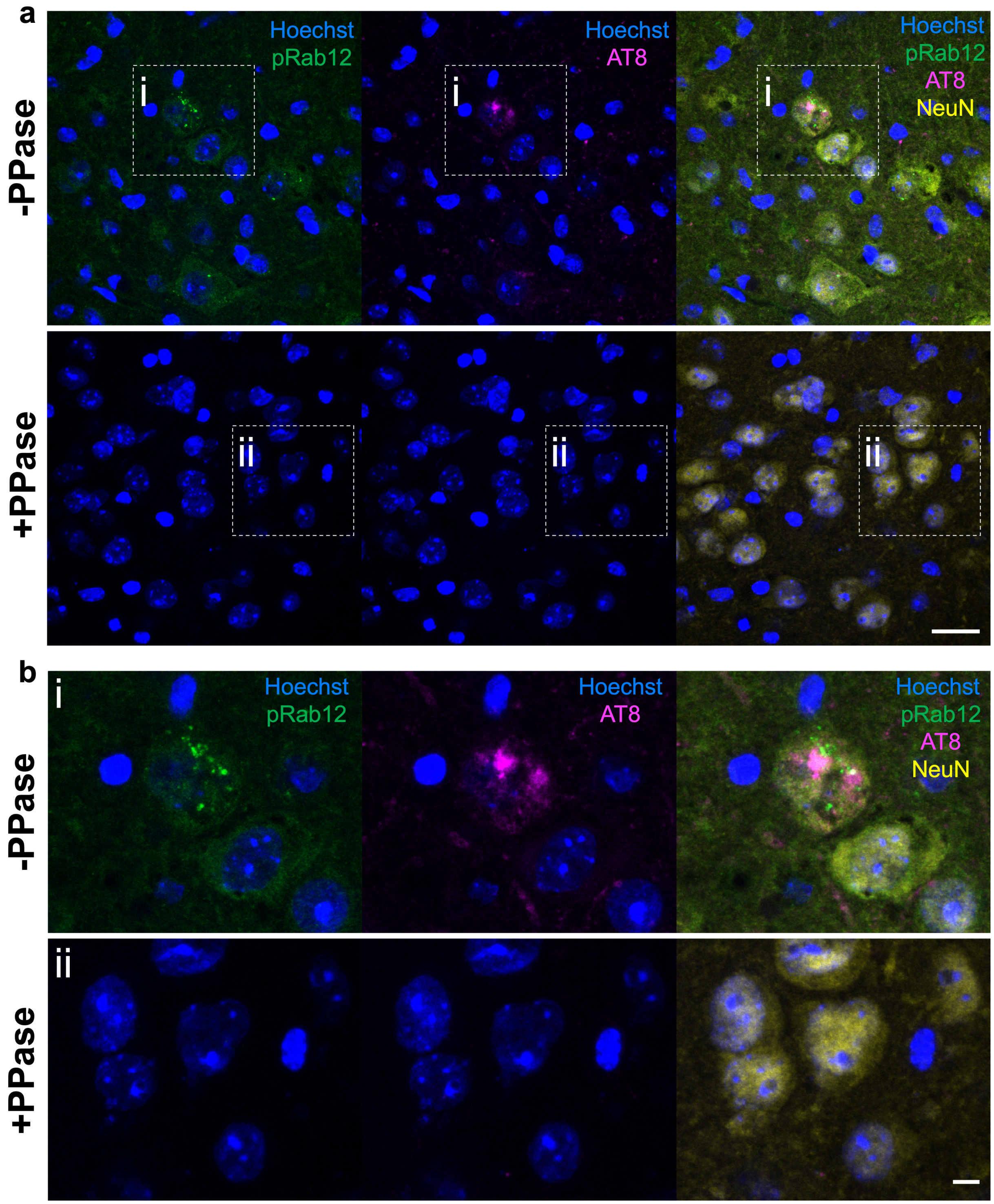


**Supplementary Fig. 11. Phosphatase (PPase) pre-treatment abolishes pS106-Rab12 and AT8 signal in aged PS19 mouse entorhinal cortex.** (**a**) Representative 63x IF images and (**b**) insets of Hoechst (blue), pS106-Rab12 (pRab12, green), AT8 (magenta), and NeuN (yellow) in aged PS19 mouse entorhinal cortex with (+) or without (-) PPase pre-treatment. Scale bar in (**a**) = 20μm, scale bar in (**b**) = 5μm.

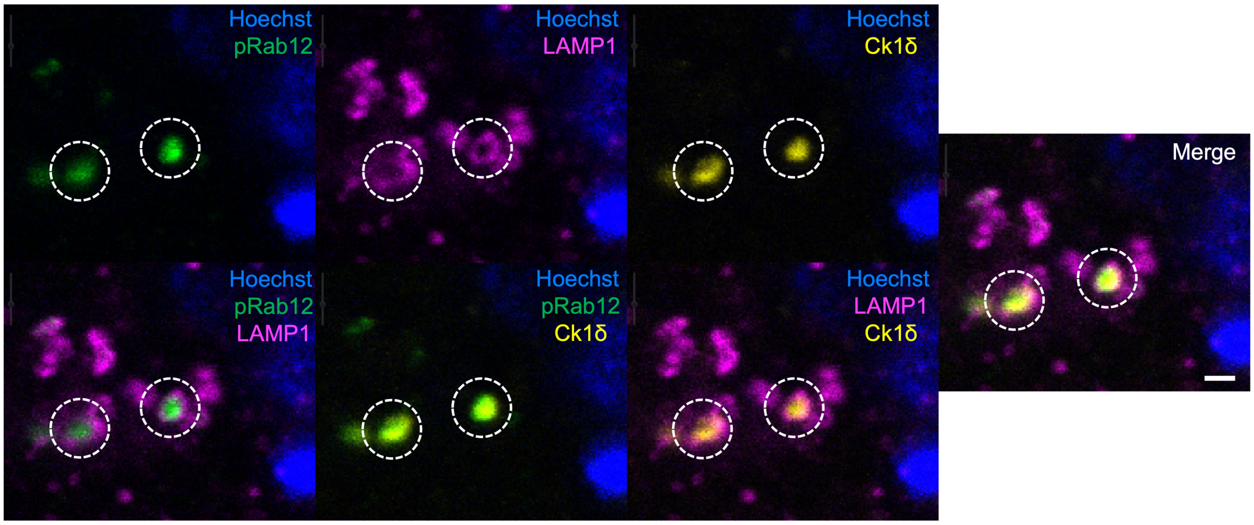


**Supplementary Fig. 12. pS106-Rab12 localizes to the GVB dense core surrounded by a LAMP1-positive membrane.** Additional representative 63x single-plane images of Hoechst (blue), pS106-Rab12 (pRab12, green), LAMP1 (magenta) and Ck1δ (yellow) in aged PS19 entorhinal cortex. Dashed circles show pS106-Rab12- and Ck1δ-positive dense GVB cores surrounded by a LAMP1-positive outer membrane. Scale bar = 1μm.

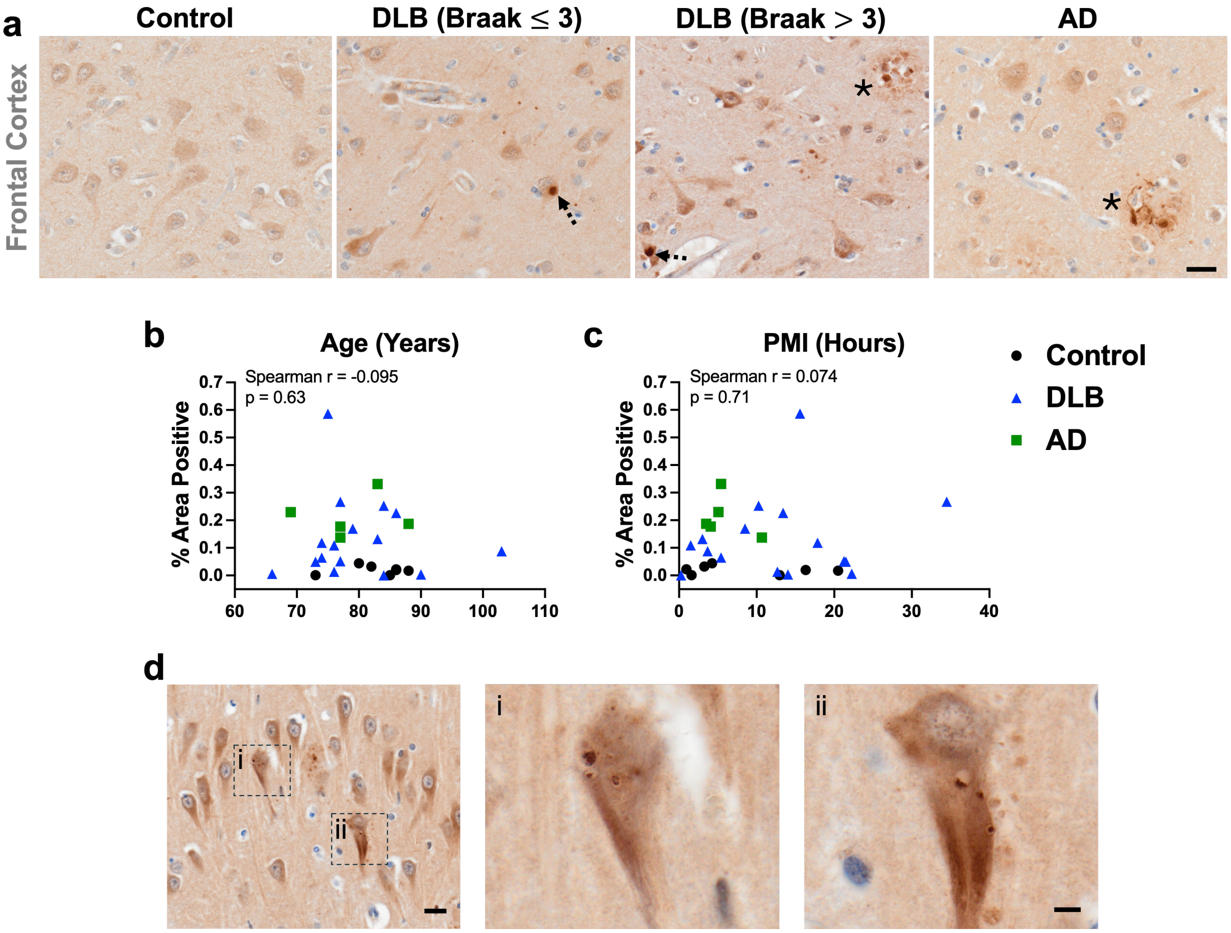


**Supplementary Fig. 13. Regional difference in pS106-Rab12-positive area, correlations of pS106-Rab12-positive area with age and PMI, and rare pS106-Rab12 GVBs alongside neurofibrillary tangles.** (**a**) Representative 60x images of pS106-Rab12 in frontal cortex of a control, DLB (Braak ≤ 3), DLB (Braak > 3), and AD case. (**b, c**) Correlations (with Spearman correlation coefficients and associated p-values) of hippocampus pS106-Rab12-positive area with (**b**) age and (**c**) PMI in control (n = 7), DLB (n = 16) and AD (n = 5) cases. (**d**) Representative 60x image and insets of an unaffected control hippocampus (Braak stage 3) showing rare pS106-Rab12-positive GVBs and neurofibrillary tangles in the same neuron. Data are presented as mean ± SEM, and each point represents an individual subject. Scale bar in (**a**) = 20μm. Scale bar in (**d**) = 20μm, inset scale bar in (**d**) = 5μm.

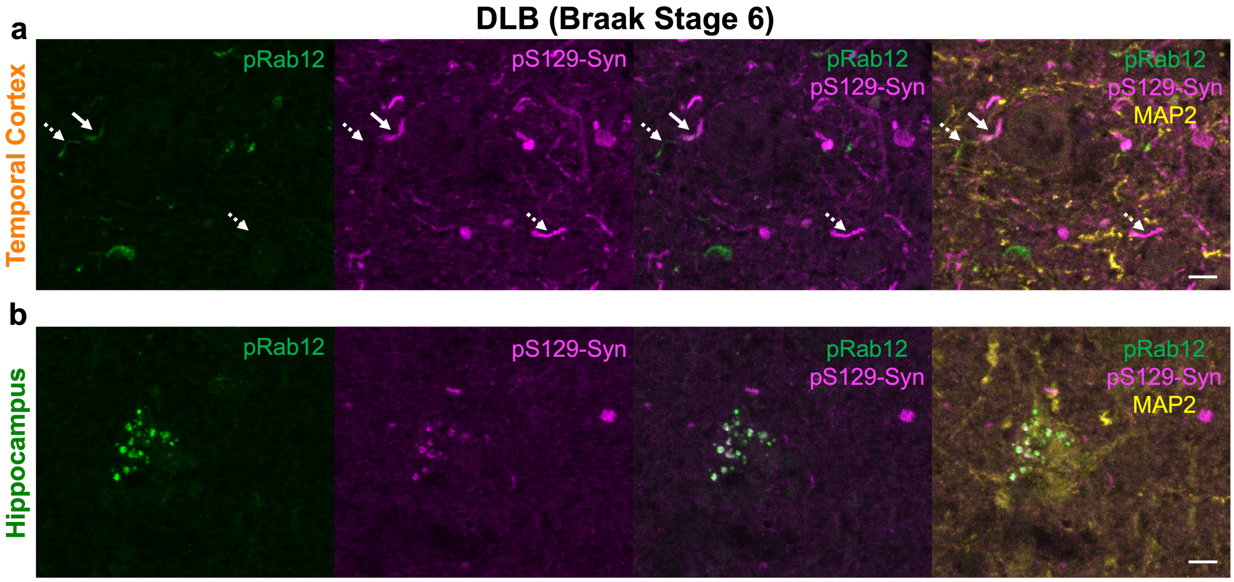


**Supplementary Fig. 14. IF reveals pS106-Rab12 labels a subset of α-synuclein pathology in DLB.** (**a**) Representative 63x IF images of pS106-Rab12 (pRab12, green), pS129-Syn (magenta), and MAP2 (yellow) in temporal cortex of a DLB (Braak tangle stage 6) case. Solid white arrows show a pRab12-positive Lewy neurite, dashed white arrows show pRab12 and pS129-Syn neurites that are not positive for the other marker. (**b**) Representative 63x IF images of pRab12 (green), pS129-Syn (magenta), and MAP2 (yellow) in hippocampus of a DLB (Braak tangle stage 6) subject showing pRab12-positive GVBs co-labeled by pS129-Syn. Scale bars in (**a, b**) = 5μm.

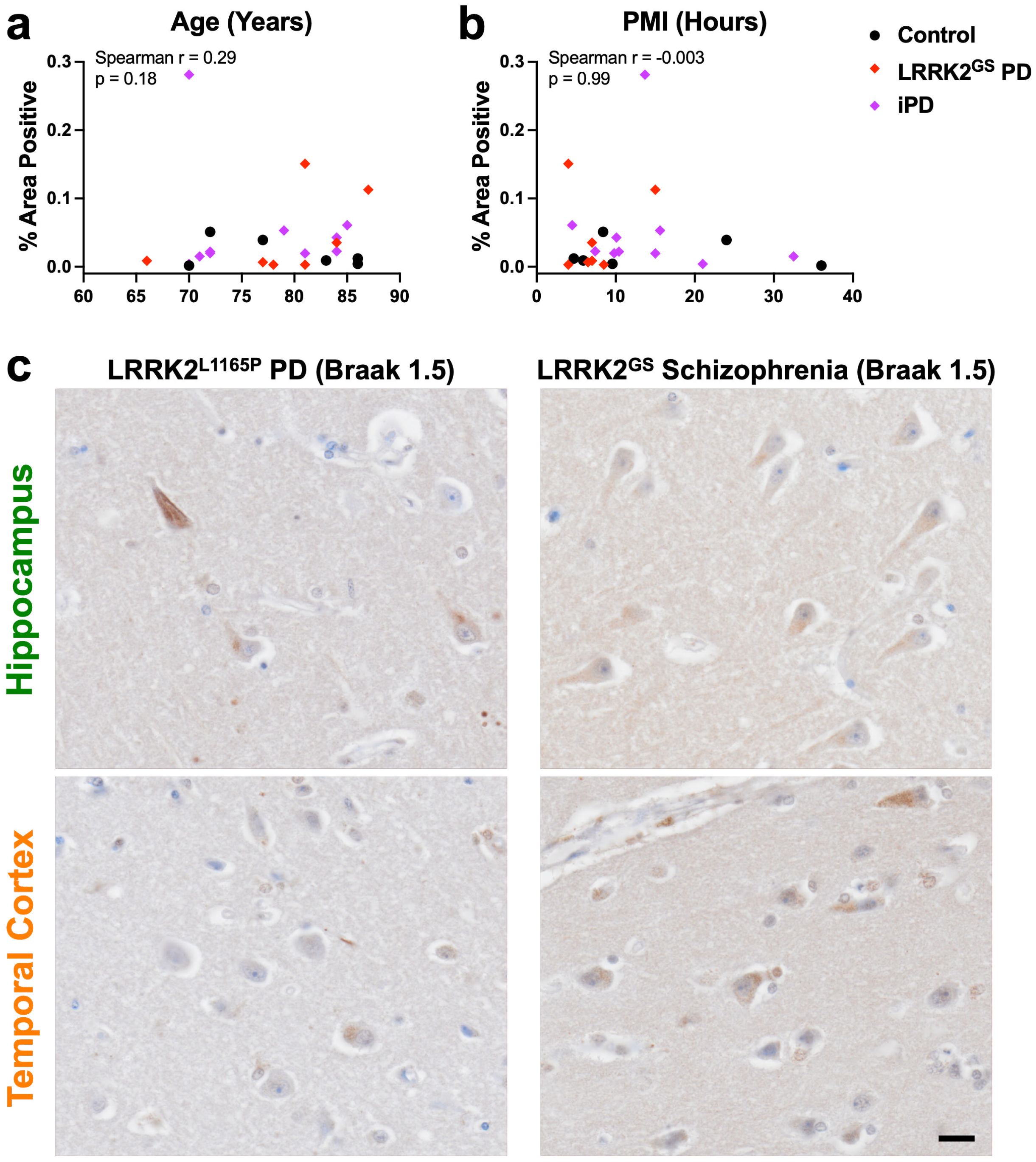


**Supplementary Fig. 15. Correlations of pS106-Rab12-positive area with age and PMI in control, LRRK2$^{GS}$ PD and iPD cases, and lack of pS106-Rab12 labeling of tau or Lewy pathology in LRRK2$^{L1165P}$ PD/LRRK2$^{GS}$ Schizophrenia.** (**a**, **b**) Correlations (with Spearman correlation coefficients and associated p-values) of hippocampus pS106-Rab12-positive area with (**a**) age and (**b**) PMI in control ($n = 6$), LRRK2$^{GS}$ PD ($n = 7$) and iPD ($n = 10$) cases. (**c**)

Representative 60x images of pS106-Rab12 labeling in hippocampus and temporal cortex of a $LRRK2^{L1165P}$ PD and a $LRRK2^{GS}$ Schizophrenia case. Each point represents an individual subject. Scale bar = 20μm.

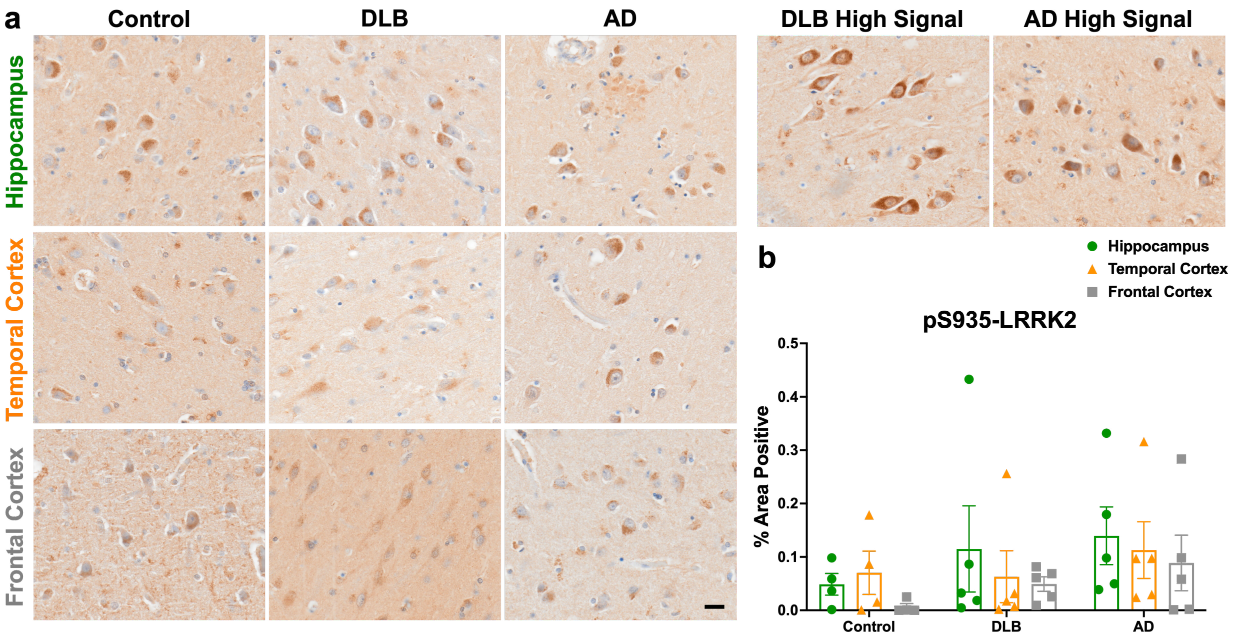


**Supplementary Fig. 16. Lack of significant difference in pS935-LRRK2 labeling in DLB and AD.** (**a**) Representative 60x images of pS935-LRRK2 labeling in hippocampus of a control, DLB and AD case. (**b**) Quantification of pS935-LRRK2-positive area in control (n = 5), DLB (n = 5) and AD cases (n = 5). Data are presented as mean ± SEM, and each point represents an individual subject. Scale bar in (**a**) = 20μm.

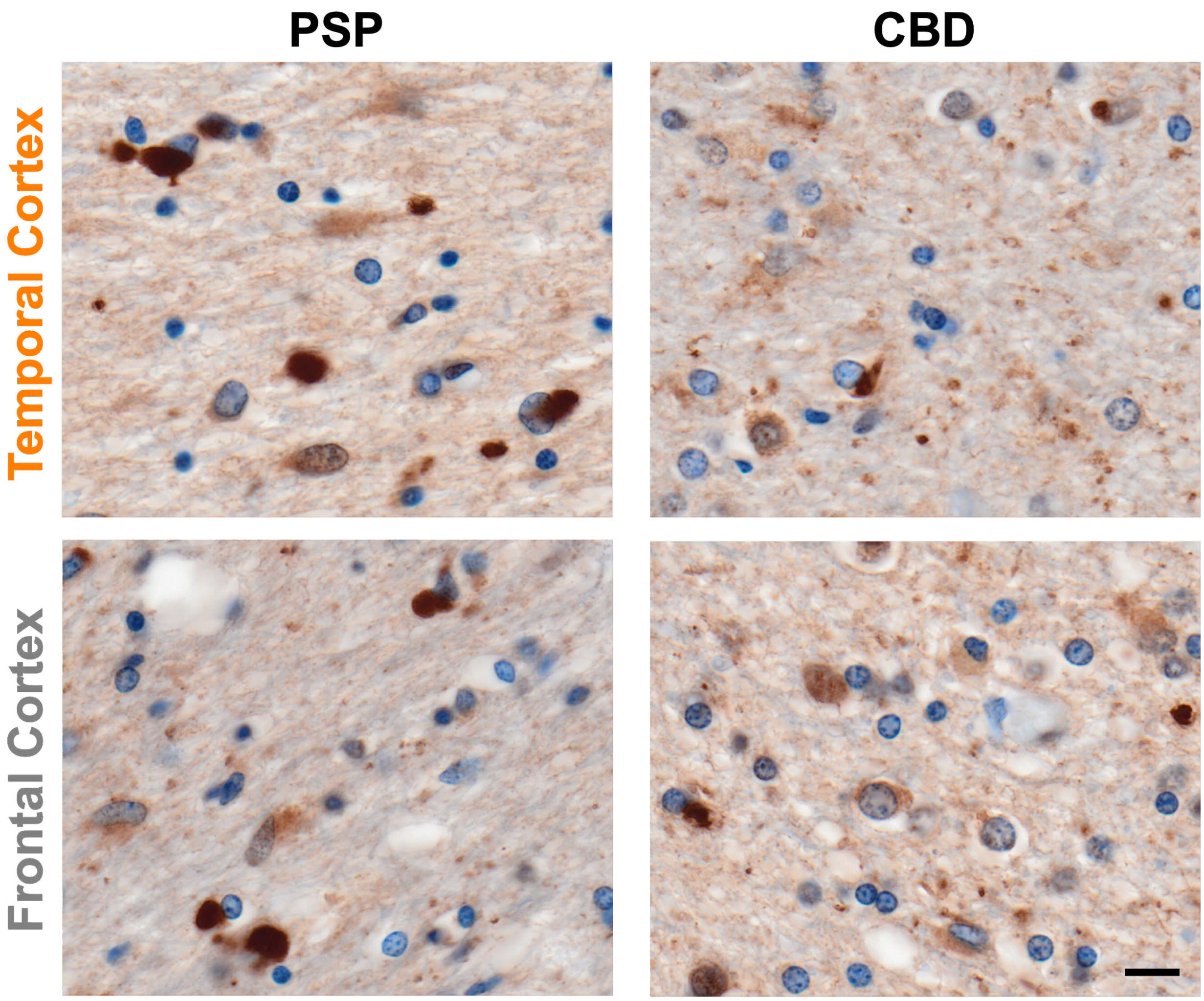


**Supplementary Fig. 17. Glial pS106-Rab12 labeling in PSP and CBD temporal cortex and frontal cortex.** Scale bar = 10µm.
